# Plasticity and Language in the Anesthetized Human Hippocampus

**DOI:** 10.1101/2025.04.09.648012

**Authors:** Kalman A. Katlowitz, Eric R. Cole, Elizabeth A. Mickiewicz, Shraddha Shah, Melissa C. Franch, Joshua Adkinson, James L. Belanger, Raissa K. Mathura, Domokos Meszéna, Matthew McGinley, William Muñoz, Garrett P. Banks, Sydney S. Cash, Chih-Wei Hsu, Angelique C. Paulk, Nicole R. Provenza, Andrew Watrous, Ziv Williams, Alica M. Goldman, Vaishnav Krishnan, Atul Maheshwari, Sarah R. Heilbronner, Robert Kim, Nuttida Rungratsameetaweemana, Benjamin Y. Hayden, Sameer A. Sheth

**Affiliations:** Department of Neurosurgery, Baylor College of Medicine, Houston, TX, USA and Neuroengineering Initiative, Rice University, Houston, TX, USA; Department of Linguistics, Rice University, Houston, TX, USA; Department of Neurology, Massachusetts General Hospital, Harvard Medical School, Boston MA USA; HUN-REN Research Centre for Natural Sciences, Budapest, Hungary; PPCU Faculty of Information Technology and Bionics, Budapest, Hungary; Department of Neuroscience, Baylor College of Medicine, Houston, TX, USA; Department of Neurosurgery, Massachusetts General Hospital, Harvard Medical School, Boston MA USA; Department of Integrative Physiology, Baylor College of Medicine, Houston, TX, USA; Department of Electrical & Computer Engineering, Rice University, Houston, TX, USA; Department of Neurology, Baylor College of Medicine, Houston, TX, USA; Department of Neurology, Cedars-Sinai Medical Center, Los Angeles CA, USA; Department of Biomedical Engineering, Columbia University, New York, NY USA

## Abstract

Consciousness is a fundamental component of cognition,^1^ but the degree to which higher-order pattern recognition relies on it remains disputed.^2,3^ Here we demonstrate the persistence of oddball discrimination, semantic processing, and online prediction in individuals under general anesthesia-induced loss of consciousness.^4,5^ Using high-density Neuropixels microelectrodes^6^ to record both single unit and local field potential neural activity in the human hippocampus while playing a series of tones to anesthetized patients, we found that hippocampal neurons and local oscillations retained some detection of oddball tones. This effect size grew over the course of the experiment (∼10 minutes), demonstrating representational plasticity. A biologically plausible recurrent neural network model showed that learning and oddball representation are an emergent property of flexible tone discrimination. Moreover, when we played language stimuli, single units and local field potentials carried information about the semantic and grammatical features of natural speech, even predicting semantic information about upcoming words. Together these results indicate that in the hippocampus, which is anatomically and functionally distant from primary sensory cortices,^7^ complex processing of sensory stimuli occurs even in the unconscious state.

## MAIN TEXT

We performed intraoperative hippocampal recordings with Neuropixels probes^6^ in six patients undergoing anterior temporal lobectomies. Recordings were conducted in the anterior body after resection of the lateral temporal cortex and prior to resection of mesial temporal structures such as parahippocampal gyrus and amygdala (**Figure 1A-G**). After a brief baseline recording, we conducted recordings during presentation of auditory stimuli composed of pure tones (3 patients) or a podcast (4 patients, **Figure 1H**). Figure 1C shows the probe in the tissue in one patient. Across all recordings, we isolated 651 units. Average firing rates were low (1.8 +/-1.1 Hz). Motion artifacts, a major challenge for human cortical Neuropixels recordings^9,10^, were markedly less conspicuous than in the cortical recordings (**Figure 1I)**, presumably due to the central location of the hippocampus, and because it is anchored by the dura of the middle fossa. Consistent with this hypothesis, the reduction in motion was especially clear when we compared respiratory and heartbeat frequency bands to a control recording performed in the cortex prior to resection (p<0.001, t-test on power between 0.1 to 3 Hz of motion trajectories between hippocampal and cortical recordings).

**Figure 1.**
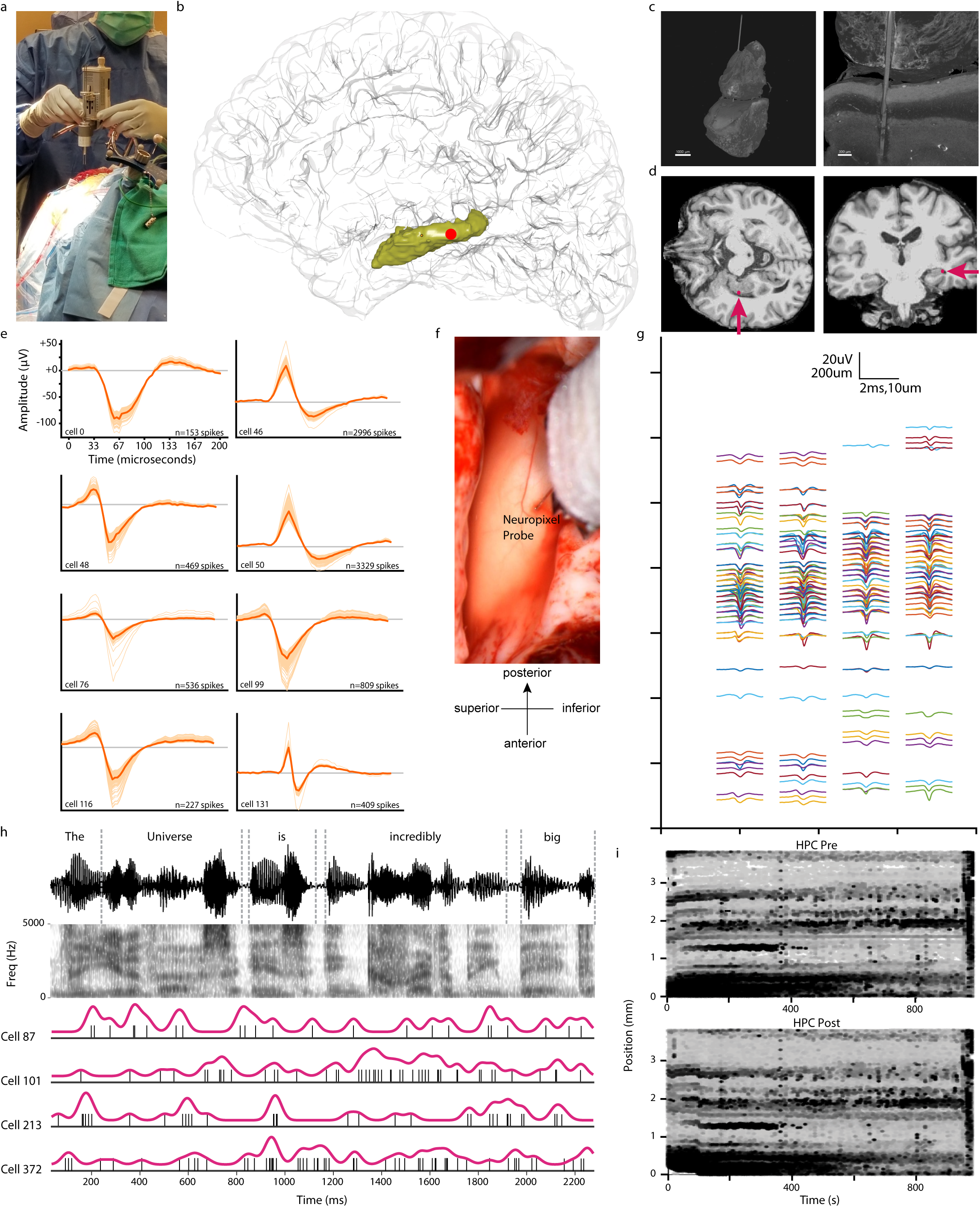
Neuropixels implantation procedure and analysis. **A.** Picture of Neuropixels insertion with robotic guidance. **B.** Probe entry site for P8 warped onto canonical brain, illustrated with a crimson dot within the hippocampus (shown in yellow). **C.** In one patient (P8), we extracted the probe along with the tissue. 3D reconstruction of microCT identifying the probe within resected hippocampal tissue (left) with coronal slice identifying the probe penetrating the hippocampus (right). Superior globule is fibrin glue adhering to the ependymal lining. **D.** Axial (left) and coronal (right) sections of a T1 MRI for P8. Crimson dot indicates probe entry site, and arrows demonstrate trajectory of probe. **E.** Example waveforms from units collected in our recording. **F.** Photograph of the hippocampal brain tissue with the inserted Neuropixels probe during intraoperative recording (middle right), with the anatomical orientation indicated below. **G.** Example waveforms from all units (n=127) within a single hippocampal recording (P5). Each unit is represented by the 15hermose waveforms at the three channels with greatest magnitude. X-axis refers to each of the four columns in the columnar arrangement of sensors. Y-axis: position along electrode shank. **H.** Analysis pipeline; we transcribed individual words with precise temporal alignment. We then computed average spike-aligned responses to words. **I.** Drift maps illustrating the low drift rates, and high stability, of Neuropixel recordings in human hippocampus. X-axis: time across session. Y-axis: depth along electrode shank. Points (black marks) indicate individual spikes. Banding pattern reflects degree of motion before (top panel) and after (bottom panel) motion correction.

### Auditory environment monitoring during anesthesia

In the oddball task^11,12^, three patients were presented with a pattern (identical 100 ms tones) interspersed with oddballs (20%, higher/lower frequency tones, **Figure 2A, Methods**). Most units (n=122/172, 70.9%, signed-rank test, α=0.05) showed tone-evoked responses (**Figure 2B**), consistent with established auditory responses in hippocampus.^13^ Tones were presented with random stimulus onset asynchrony (1-3 seconds). Across all units, response latencies showed a biphasic temporal dynamic (Gaussian Mixture Model fit via Expectation Maximization, **Figure 2B and 2C**). Units encoded tone identity (p<0.001, generalized linear mixed effects, GLME, see **Methods**; n=39/172, 22.7% of units, rank sum test, α=0.05, **Figure 2D**). There was no difference in the proportion of neurons that significantly responded to each oddball type (p=0.184; N=172 units; chi-squared goodness of fit test), thus tone discrimination effects reflect a balanced shift in the population response rather than a preferred acoustic feature of one of the tones.

**Figure 2.**
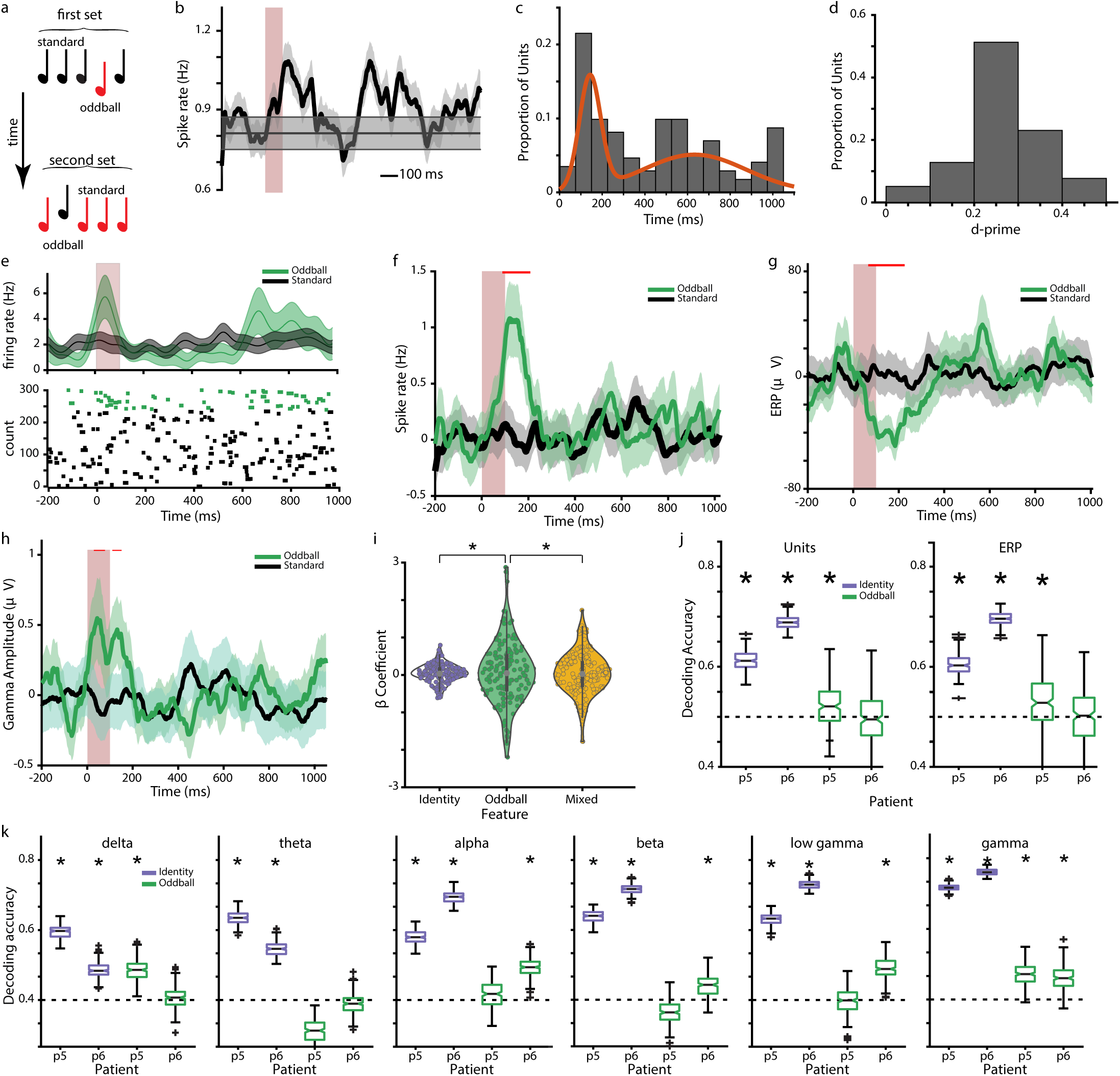
Oddball responses in the anesthetized human hippocampus. **A.** Schematic of the auditory oddball task. Each trial consisted of interleaved pure tones and oddballs. The oddball and standard were interchanged after 150 tones. **B.** Mean response to tone onset, averaged across all units (n=172 units, 3 patients). Vertical red bar: tone presentation (100 ms), horizontal grey bar: baseline +/- standard error of the mean (SEM). **C.** Distribution of the tone response onset latencies across all units. A mixed Gaussian model fit to the distribution is shown in orange. **D.** Distribution of d-prime values for units selective for tone identity. **E.** Example unit selective for oddballs. Top: Average response (spike rate, Hz) to oddball (green) and standard (black), shading: SEM, red bar: tone presentation. Bottom: trial-wise spike raster plot, color-coded as oddball (green) and standard (black). **F.** Average neuronal response across all units (n=150 units to oddball (green) vs. standard (black) tones. Red lines indicate periods of significant difference. **G.** Average ERP (μV) across 10 channels from each patient. **H.** Average gamma band amplitude (μV) across 10 channels from each patient. **I.** Violin plot showing the distribution of β coefficients obtained from a linear regression run per unit, to determine modulation as a function of tone identity (purple) oddball identity (green), and an interaction/mixed term (yellow). Asterisks: statistical significance of difference in absolute amplitude. **J.** Box and whisker plot of decoding accuracy for tone identity (purple) and oddball identity (green), obtained using a Support Vector Machine (SVM) decoder for P5 and P6, across units (left) and ERP (right) after tone presentation. **K.** SVM decoding accuracy for tone identity (purple) and oddball identity (green) for both patients, across the population of recorded channels within each pre-defined frequency band. * denotes p-value <0.001.

We next examined the representation of stimulus features. For two patients, we balanced tone identity and oddball status (**Figure 2A**, n=150 units). At the single unit (**Figure 2E**) and population (**Figure 2F**) levels, neuronal responses differentiated standard from oddball tones (p<0.0001, GLME). This divergence was most notable within the first 300 ms (24.7%, n=37/150 of units, signaled oddballs. Subsequent analyses focused on this epoch segment. Note that we use a symmetric non-causal Gaussian filter, which equally weights past and future time points; the small amount of responding before the tone is an artifact of this filter. Moreover, because excitatory and inhibitory tone-evoked responses tended to cancel each other out, there is no visible peak in the response for the standard tone. Local field potentials (LFPs) also showed oddball-evoked responses: a negative deflection in the evoked response potential (ERP, **Figure 2G**) and an increase in gamma amplitude (**Figure 2H**).

We modeled evoked responses for all units as a function of tone identity, context (standard vs. oddball), and their interaction. We observed comparable encoding for all terms: tone encoding (p<0.0001, GLME; 29.3% of units significant), oddball encoding (p<0.0001, GLME; 24.7% of units significant), and interaction (p<0.01, GLME; 22.7% of units significant). The absolute values of the betas for the oddball term were greater than the corresponding tone and mixed selectivity terms (paired t-test on absolute values, p<0.0001 for both, **Figure 2I**). Similar proportions of units showed a significant oddball effect (n=43) in the two patients (37/127, 29.1% and 6/23, 26.1%, p=0.8, 11^2^ test). Mean broadband LFP power and gamma band amplitude also demonstrated tone, oddball, and mixed selectivity at similar rates across channels but reduced significance (broadband LFP: p = 0.224, p = 0.038, and p = 0.0186, LME; 40.9%, 47.2%, and 46.0% of channels. gamma: p = 6.18e-4, p = 0.024, and p = 2.80e-4, LME; 20.1%, 17.6%, and 18.7% of channels, respectively). Tone detection effects remain robust at lower significance levels of p<0.01 (55.2% of neurons) and p<0.001 (28.5%), as to tone-selective effects (p<0.01: 12.2%; p<0.001: 4.1%). Modeled oddball detection effects remain robust at p<0.01 (14.7%) and p<0.001 (9.3%), and tone/oddball interaction effects remained robust at p<0.01 (16.7%) and p<0.001 (10.0%).

We used a 10-fold cross-validated support vector machine (SVM) to decode stimulus features on a trial-by-trial level across the population. Tone identity was robustly represented in both patients across units and LFP, with accuracy ranging between 0.61 and 0.70 (p<0.001 for all, t-test, **Figure 2J**). Oddball identity was decodable at above chance for the two patients combined (p<0.05), albeit not in p6 alone. We found that tone identity could be decoded across all tested frequency bands (range: 0.52 to 0.79, p<0.001, t-test). Oddball identity was decodable in some frequency bands for both patients (p<0.001 for delta and high gamma bands for p5; all bands except delta and theta for p6). Thus, tone and oddball information is present in both single unit and LFP, with oddball signals having substantially weaker strength than tone identity.

High gamma phase-amplitude coupling reliably detected tone and oddball stimuli: 26.6% of channels demonstrated significant tone encoding, 21.96% of channels demonstrated significant oddball encoding, and 24.07% of channels demonstrated an interaction (p < 0.05). Leveraging SVM decoding, high gamma phase-amplitude coupling predicted tone identity (p5: tone accuracy = 0.553; p6: tone accuracy = 0.542), but could not decode oddballs. Low gamma phase-amplitude coupling did not significantly encode tones, oddballs, or their interaction. Low gamma phase-amplitude coupling could weakly predict both tone identity and oddball status in p6 (p6: tone accuracy = 0.505, p < 0.001; oddball accuracy = 0.527, p < 0.005), but not p5 (p > 0.05 for both tone and oddball).

### Signatures of plasticity in the unconscious state

We next examined the temporal evolution of the oddball response. In oddball-selective units (n=43), we found that oddball response grew more distinct (as inferred from decodability) over the course of the experiment (∼10 minutes, example unit: **Figure 3A)**. Splitting our task into first and second halves (for each block), we found a significant increase in oddball encoding for both patients (P5: p=0.01, P6: p<0.001, t-test, **Figure 3C**). We also observed a decrease in tone identity encoding from the first to the second half of each block, raising the possibility of compensatory mechanisms (p<0.001, t-test, **Figure 3B**). Using a 50-stimulus sliding window, we found a continuous increase in oddball decoding accuracy across the 10-minute duration of the experiment (p<0.001, Pearson’s correlation, **Figure 3D, purple**). This increase in oddball performance was accompanied by a decrease in tone encoding (p<0.0001, **Figure 3D, green**).^15^ There was a significant negative correlation between the evolution of trajectories of oddball and tone decoding accuracy (r = -0.23, p<0.03, Pearson’s test), consistent with the hypothesis that the neural population was sacrificing tone responses for the sake of oddball representations over the course of the experiment.^16^ Phase-amplitude coupling demonstrated a significant difference in tone decoding accuracy between first half and second half of the oddball experiment (t-test; p<0.0001), in both patients. Oddball decoding only significantly increased in the second half of the oddball task for p6 (p<0.0001), not p5. Spike-frequency coupling showed a significant increase in both tone and oddball decoding between task halves, and in both patients (p<0.0001).

**Figure 3:**
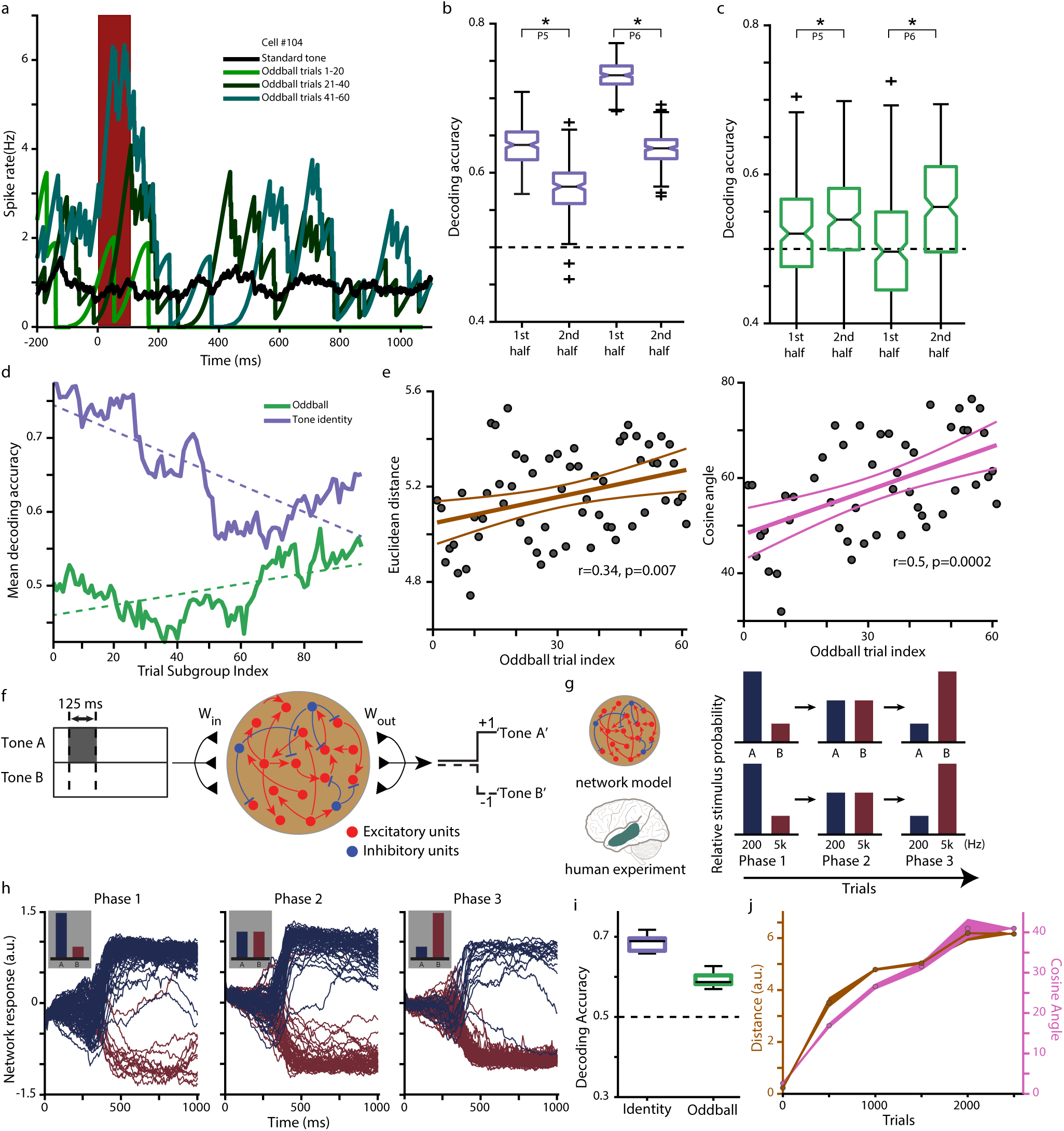
Evolution of the oddball representation across the neuronal population in experimental data and an RNN model. **A.** Responses to tones as a function of oddball identity and index in an example unit. Red bar indicates tone presentation. **B.** Accuracy of tone identity decoding across the neuronal population for patients P5 and P6, for the first half of each block (left) or second half of each block (right), combined across both blocks. Statistically significant differences indicated with an asterisk **C.** Similar to **B** but for oddball identity. **D.** Decoding accuracy as a function of trial position across both patients (n=43 oddball-responsive units). Each point represents SVM accuracy within a set of 50 trials starting at the index location. Decoding accuracy for tone identity shown in purple, and for oddball identity shown in green, along with line fits shown as dashed lines in purple and green respectively. **E.** Euclidean distance (left) and cosine angle (right) between standard and oddball neuronal population response vectors, computed for each oddball trial. Each datapoint (in grey) indicates the value of the Euclidean distance (left) and cosine angle (right) per trial, with the lines showing a linear fit with 95% confidence intervals **F.** Schematic of the RNN model trained to differentiate between two different tone frequencies, indicated as Tone A and Tone B. **G.** Training paradigm for the RNN as compared to the human experiment. **H.** Network response across trials for a single example RNN model unit for the three training phases. **I.** Decoding accuracy of RNN for tone identity (purple, left) and oddball identity (green, right). **J.** Evolution of Euclidean distance (brown) and cosine angle (pink) between oddball trials and the average standard trial across the RNN population. Shading represents SEM across 10 runs.

We created neural vectors of the average standard tone as well as each individual oddball trial (43-dimensional vectors composed of the mean response of the oddball units). We found a gradual divergence in Euclidean distance between standard and oddball vectors over the course of the session (r=0.34, p < 0.0001, LME; **Figure 3E, left**). Discriminability was even stronger when considering cosine angle, indicating the effect is not merely a consequence of a response gain in oddball cells (r=0.5, p<0.0001, LME; **Figure 3E, right**). These effects were mostly consistent for individual patients (P5 distance: r=0.25; p=0.056, angle: r=0.43 p=0.002; P6 distance: r=0.32, p=0.012, angle: r=0.48; p=0.0002). These results indicate that the hippocampus does not simply improve encoding using gain modulation;^17^ instead, oddball responses reflect a rotation of the neural population vector in a high dimensional space, meaning that neural plasticity alters the shape of the neural response manifold.^18^ Thus, complex reshaping of responses can occur even under general anesthesia.^19,20^ Further analyses of the information encoded in LFP, including band-limited analysis and aperiodic slope can be found in Supplementary Figures S1-S5. Additional analyses relating to cell-type encodings can be found in Supplementary Figures S6-S7.

To gain further mechanistic insight at the level of individual units, we turned to a continuous-rate recurrent neural network (RNN) trained to perform a signal-detection task similar to the task used for the human Neuropixels data (**Figure 3F**).^21^ The network model underwent three stages of training, simulating the different contexts used in the experimental data, with the prevalence of specific tones varied at each stage (**Figure 3G, H, Methods**). To simulate the two auditory tones, the model received Gaussian white noise across two input channels, each corresponding to one of the two tones (Tone A and Tone B). On each trial, a transient +1.0 bias was added to one of the two channels during a brief stimulus window, indicating the presence of the corresponding tone.

Tone A was presented to the network in 80% of the trials, followed by a washout period, and then a stage with probabilities reversed relative to the first. By the end of training (range of 1400 to 2600 trials) the model was able to differentiate tone identities (**Figure 3H**). Notably, despite being explicitly trained only on tone identity discrimination, the model was able to perform not only identity discrimination (tone identity, p<0.005, signed Wilcoxon test vs. shuffled data) but also context classification (oddball identity, p<0.005, signed Wilcoxon test vs. shuffled data, **Figure 3I**) via linear SVM decoding (see **Methods**). The model also recreated the pattern observed in the Euclidean and vector angle distance between standard and oddball representations (**Figure 3J)**, suggesting that the divergence of representations can be due to local computations rather than inherited from other networks. Note that although the standard tone occurs in 80% of trials, the network was trained to make a binary choice between two alternatives. Therefore, the theoretical chance performance remains 50%.

Rate-based RNNs provide a tractable framework for investigating how E-I interactions give raise to these mechanisms. To this end, we used an E-I RNN model to reflect the known composition of cortical circuits, which are predominantly excitatory (∼80%) with a smaller proportion of inhibitory neurons (∼20%). To more fully leverage the EI structure of the model, we conducted systematic lesioning analyses in which we selectively lesioned each of the four recurrent connection subtypes (E->I, E->E, I->E, I->I) by setting the corresponding weights to zero. We then re-computed SVM decoding performance for both tone identity and oddball context.

Systematic lesioning of each of the four recurrent synaptic connection types (E→E, E→I, I→E, I→I) revealed that inhibitory connections are essential for encoding both tone and oddball categories (see Supplementary Figure S8). Lesioning inhibitory-to-excitatory (I->E) and inhibitory-to-inhibitory (I->I) connections led to the most pronounced drop in decoding accuracy for both tone identity and oddball context. These findings indicate that inhibitory feedback, both directly onto excitatory neurons and within inhibitory populations, plays a critical role in shaping population-level representations. Inhibitory connections were important for encoding both tone identity and oddball context.

### Unconscious encoding of semantics and grammar in the hippocampus

We next tested whether the unconscious hippocampus could perform even higher order functions associated with parsing semantic and syntactic features of natural speech. In four participants (P6, P8, P9, and P11), we recorded neural activity while playing 10-20 minutes of podcast episodes (see **Methods**^22^). We aligned neural activity to word onset and offset (n=3024 words for P6; n=1565 words for P8 and P9; n=962 words for P11), and computed word-evoked neural responses (example unit, average response to all words presented, **Figure 4A**).

**Figure 4.**
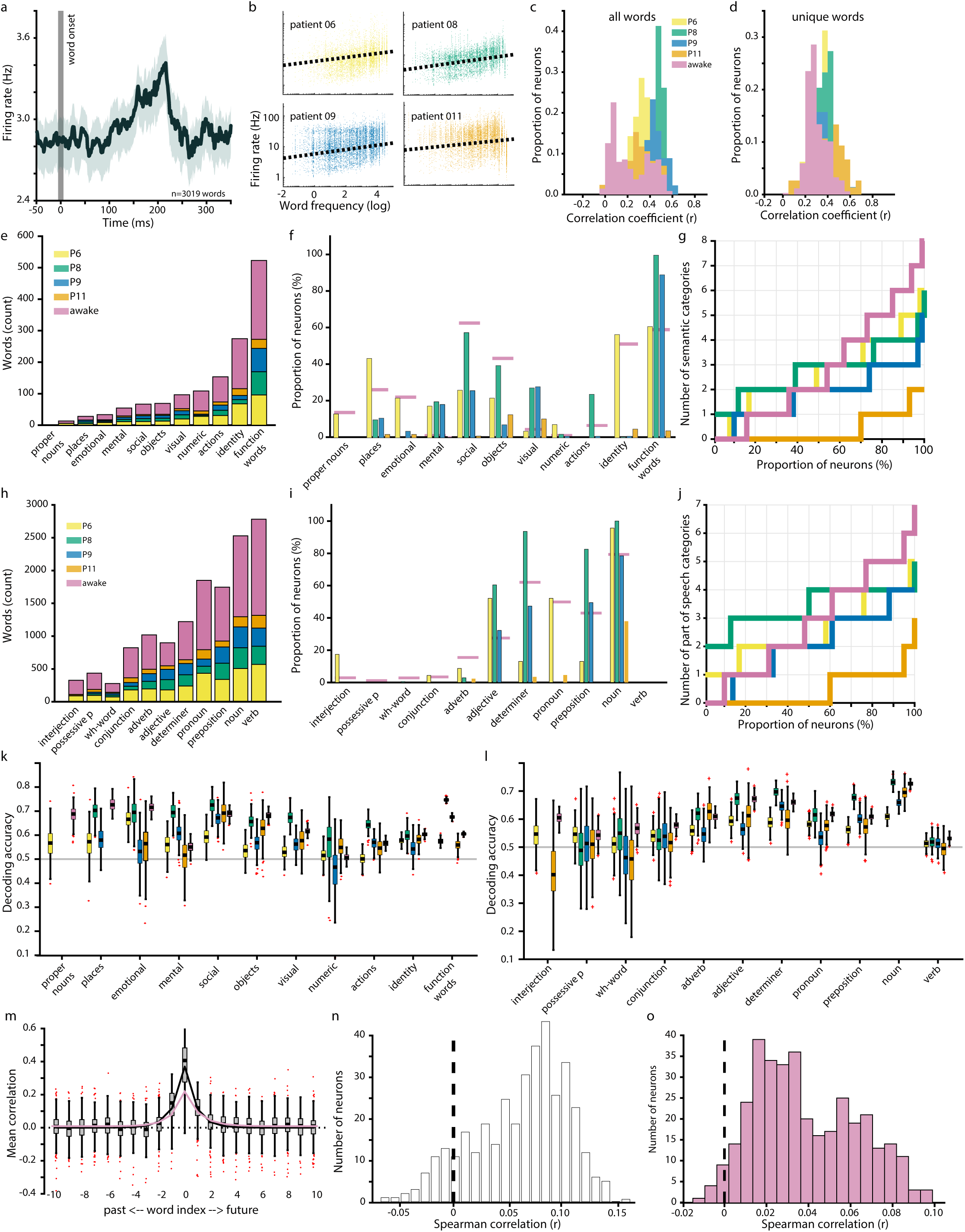
Language responses in the anesthetized human hippocampus. **A.** Average response (spike rate, Hz) of an example unit across 3019 words, aligned to word onset (vertical dashed line). Shaded area: SEM. **B.** Neuronal responses as a function of word frequency shown for each neuron. Each data point corresponds to a neuron and word, color-coded according to patient. Dashed line: linear fit for that patient. **C-D.** Distribution of Pearson’s correlation coefficient for predicted vs. actual firing rates for all I and unique words (**D**), data shown per patient. All correlations are greater than zero, indicating that the models are better than chance at fitting neural responses. **E.** Distribution of words within semantic categories sorted by frequency within the stimulus set; per patient. **F.** Percentage of units selective for each semantic category, compared to all other semantic categories, per patient. **G.** Number of categories decoded by individual units, per patient. Neurons generally participate in coding of multiple categories. **H-J.** Similar to E-G but for part of speech categories. **K-L.** Decoding accuracy of a support vector machine (SVM) across units for semantic (**K**) and part of speech (**L**) categories, shown per patient. Dashed lines represent chance; pluses indicate outliers. **M.** Correlation coefficients for predicted vs. actual firing rates as a function of word Index. Index=0: current word. Negative indices indicate past words; positive indices indicate future words. **N and O.** Histogram of the correlation coefficient (Spearman) for each neuron between surprisal and firing rate in anesthetized (N) and awake (O) patients. Overall, firing rates were higher for surprising words.

Given the oddball effects described above, we first hypothesized that the brain would respond differentially based on word lexical frequency, which we defined using a standard database.^23^ We found a statistically significant correlation between firing rate and word frequency in all four patients individually (**Figure 4B and Figure S9**). Specifically, we find a significant positive correlation in the single unit activity (mean r=0.48+/- 0.06, Spearman’s correlation, α<0.05) and across patients (p<0.0001, GLME) and a modest but significantly negative correlation in all six bands of the LFP (range -0.08 in Beta band to -0.02 in Delta band, p<0.001 for all). Notably, lexical duration is also significantly encoded (p<0.0001, GLME). To address possible confounds between word duration and frequency, we reran the unit analysis with subsets of words within a limited duration range, i.e. 0-200 ms, 200-400 ms, 400-600 ms, and consistently observed a positive correlation (p<0.001 for combined units in each patient separately). Additionally, a linear model that incorporated both logarithmic word duration and logarithmic word frequency still found significance in word frequency as a predictor of firing rate (p<0.001, in each patient separately, t-test on coefficients). This correlation could not be solely explained by difficulty in lexical access, as there was also a consistent relationship of the neural responses with the relative surprise of each word (see below).

These results suggest that the unconscious hippocampus has access to the semantic information conveyed by each word. To explicitly test this possibility, we regressed the firing rates of each neuron against the semantic embeddings of each word that demonstrated a response (see **Methods**).^22,25,26^ In semantic embedding space, similar words (e.g. ‘dog’ and ‘cat’) are closer (Euclidean distance, d=1.8) than dissimilar words (e.g. ‘dog’ and ‘pen’, d=2.5). Using 10-fold cross-validation, we found that the RMSE of a linear model outperformed shuffled data in all units (α=0.05, one tailed t-test on real versus shuffled RMSE, **Figure 4C**), with an average correlation between true and predicted firing rates of 0.397 +/- 0.007 (n=368 units).

Overall, these results parallel those obtained from a separate cohort of awake patients who performed a similar task in the EMU with single units recorded on microwire electrodes (Franch et al., 2025). Specifically, in awake patients, the mean correlation was slightly lower, at 0.226 +/- 0.009, (356 units across 10 patients). However, given that conversational English has many words that are repeated these results could theoretically be confounded by the fact that cells had consistent responses to words, perhaps even matching acoustic features. To show that units generalize across word embeddings, we re-ran the analysis using only unique words (n=743, 571, 571, and 329 words for the four patients, respectively). We found a significant result in 75.4% of the recorded units (251/333 units with at least 50 words that had a non-zero response), with an average correlation of r=0.207 +/-0.05, **Figure 4D**). These numbers were again comparable to the numbers observed in the awake cohort (72.3% of units; 246/336; mean r=0.134 +/- 0.005). In other words, it is possible to predict the firing rate of units to a given word based on responses to other words by leveraging their similarities in semantic space,^27^ demonstrating that the unconscious hippocampus has access to abstract semantic relationships between words.

We then analyzed the representation of word features. We placed each word into one of 12 semantic categories (**Figure 4E)**.^22^ Nearly all units (85.6%, n=321/375) showed some form of semantic category selectivity (α=0.05, Kruskal Wallis test for any difference between semantic categories, **Figure 4F**). This number is similar to the corresponding number in awake patients (76.1%, n=271/356). Units were selective for multiple semantic categories, consistent with our previously reported findings in awake patients (rank-sum test, corrected for multiple comparisons, α<0.05).^22^ Specifically, 239/375 (63.7%) of units discriminated at least two (of twelve) categories; 54/375 (14.4%) discriminates at least four (**Figure 4G**), median of three categories per neuron. The corresponding numbers in awake patients were 232/356 and 139/356, respectively. We found that 298/375 units carried information about part of speech^28^ (α=0.05, Kruskal Wallis test, **Figure 4H**); nearly identical numbers for awake patients: 298/356. Again, there was broad representation of different categories (**Figure 4I**). Most units (80.0%) distinguished nouns from non-nouns, but none distinguished verbs from non-verbs; these numbers were comparable in awake patients (79.5% and 0%). Overall, the median number of part of speech categories represented was four (out of 11, **Figure 4J**), with 259/375 (69.1%) units discriminating at least two categories and 106/375 (28.3%) discriminating at least four.

These numbers were similar for awake patients: 251/356 (70.5%) units discriminated at least 2 categories; 141/356 (39.6%) units discriminated at least 4 categories. Interestingly, we found a modest, but significant correlation between the number of semantic categories and the number of part of speech categories represented across neurons (r=0.34, p<0.001 Spearman’s correlation; awake: r=0.24; p<0.01) suggesting that language responsive neurons can represent multiple features. Word frequency encoding effects are robust using p<0.01 (98.1%; awakes: 86.8%) and at p<0.001 (97.1%; awakes: 79.8%). Semantic encoding proportions are also robust using p<0.01 (77.9% and 68.5%) and p<0.001 (62.1% and 52.0%,). Likewise, part of speech: p<0.01 (71.5% and 62.1%) and p<0.001 (74.4% and 62.6%).

We next examined hippocampal decoding ability on a word-by-word basis. We used SVM to compare each category against all others. We found that all categories in both semantics and part of speech were decodable at the level of individual words (p<0.001 vs. shuffled data, **Figure 4K and L**). (Note that chance is 0.5 because we subsampled the majority class such that positive and negative data had equal frequency). Semantic categories had a higher average classification accuracy (60.5 +/- 4.0%) than part of speech categories (56.5 +/- 5.3%, p=0.03, t-test). Thus, both semantic and syntactic information is decodable in real time within the anesthetized hippocampus.

We next asked whether the anesthetized hippocampus could represent recent or upcoming words.^30^ We reran our linear regressions predicting previous and upcoming words (using held-out data to prevent overfitting; **Methods**). We found that neural responses corresponded not only to semantic features of prior words (**Figure 4M**, negative indices), which could be due to short term memory^31^, but also to the semantics of future words^32^ (**Figure 4M**, positive indices). Future words were decoded nearly as well as past words (across all four patients: β_0_=0.370, τ_future=0.840; τ_past=0.868). These numbers are comparable to those for awake patients (β_0_=0.227; τ_future=1.081; τ_past=0.895); indeed, they do not differ significantly (p>0.05 in all cases in the range +1 to +5). Thus, not only is recent speech actively maintained, but encoding of words is contextualized such that we can decode future words from these encodings; this type of contextualization is crucial to speech comprehension.^33^ (These results do not necessarily imply active prediction beyond contextualization^34^). Moreover, firing rates were modulated by surprisal, a metric that quantifies the probability of each word as a function of prior words,^24^ in many units (r=0.06 +/- 0.0023, 246/375 units significant, **Figure 4N and 4O**). Similar results were observed in awake patients (r=0.0386 +/- 0.0013, 245/356 neurons significant). Surprisal effects are also robust at p<0.01 and p<0.001 in anesthetized (56.8% and 40.3%, respectively) and in awake patients (56.2% and 43.0%, respectively).

Further analyses of the LFP are found in Supplementary Figures S10-S13. In short, we found that semantics were encoded in all six bands (Figure S10), although less faithfully than in unit data (Figure 4C). We also found encoding of semantic category in all bands (Figure S11), and, again, less strongly than with the single units (Figure 4F-G). Likewise, we found significant encoding of part of speech in all frequency bands (Figure S12), although less strongly than semantic category (Figure 4I-J). An SVM approach was able to classify part of speech and semantic category (Figure S13) about as well as the single units in all bands (Figure 4K-L). Aperiodic slope outperformed individual spectral features in predicting semantic embeddings (mean r = 0.32 +/- 0.17, rank sum test, p<0.001 for all bands) and robustly encoded multiple semantic and POS categories (62.5% and 25.0% encoded at least 8 categories, respectively).

We also found robust phase-amplitude coupling results for the language data. Theta-low gamma phase-amplitude coupling outperformed spectral features in predicting semantic embeddings (mean r = 0.170 +/- 0.13, rank sum test, p<0.001 for all bands) and similarly encoded multiple semantic and POS categories (74.5% and 73.0% encoded at least 8 categories, respectively). Phase-amplitude coupling using the high gamma band (70-150 Hz) showed comparable semantic embedding prediction (mean r = 0.166 +/- 0.13) and improved encoding of semantic and POS categories (99.3% and 98.7% encoded at least 8 categories, respectively).

However, spike-frequency coupling in the theta band did not significantly encode semantic embeddings (mean r=0.002 +/- 0.004) or categories (7% and 2% of units encoded one semantic and POS category, respectively; none encoded 2 or more). Together, these results attest to the robust recoverability of linguistic information in the anesthetized brain.

## DISCUSSION

We identified neural signatures of plasticity and semantic processing in the anesthetized human hippocampus. Our findings do not have obvious explanations based solely on low-level sensory responses. For example, the long and slow increase in oddball detection over the course of 10 minutes is unlikely to reflect adaptation or repetition suppression, which both take place on shorter timescales. Likewise, the representation of semantic features in speech listening requires specific processing of acoustic information. We therefore show that within anesthetic-induced unconsciousness some high-level process of sensory integration is preserved, suggesting that it is consolidation that is compromised. These results provide a potential explanation for previous reports of post-anesthesia implicit recall, which would depend on sensory processing and plasticity processes (REF).

Broadly speaking, these results confirm ideas previously developed in animal and human studies showing preserved neural responses to sensory stimuli, including oddball stimuli, during anesthesia (REF), and extend them to include (i) change over the timescale of several minutes, a timescale usually associated with wakeful learning, (iii) linkage to a plausible biocomputational model that avoids the types of executive control that are presumably diminished during anesthesia, (iii) availability of language information beyond the level of auditory processing. The present results complement our past studies, which were performed using awake patients in the epilepsy monitoring unit (EMU), on hippocampus neuron-level representation of lexical semantics^35^ and semantic contexualization^36^. The present results suggest that awake-like semantic responses, and at least some contextualization, can occur in the absence of conscious awareness. That in turn raises the question of how far linguistic processing can go in the absence of awareness^37^.

Some aspects of the local field potential correlate with single unit activity, although this relationship may fluctuate^38^. Much evidence suggests that the gamma band, in particular, may align closely with single units^39,41^. We find that the gamma band is generally aligned with neural activity, although not perfectly so, and that other bands are sometimes also aligned. However, failure to achieve significant effects may reflect an insufficiency of data; therefore, it is difficult to draw firm negative conclusions from observed differences between unit and LFP responses. In any case, our results do indicate that LFP activity, especially in the gamma band, can be a proxy for single units, although it may have other functional roles as well.

While the hippocampus is not a well-known part of the core language network, it has a well-established role in language^35,42–44^, including in recent single unit recording studies^35,36,45^. In particular, its known functions echo semantic contextualization: (1) it is associated with mapping functions related to temporal position encoding^46,47^, (2) it integrates information across multiple modalities and uses that information to contextualize representations in a variety of domains^48^, (3) and it is closely associated with the processes of prediction that drive contextualization^49,50^. Finally, (4) it has a prominent role in memory^51,52^. Some of the reason for the absence of the hippocampus in classical language models is negative evidence – it is difficult to image using fMRI and lesions isolated to the hippocampus are rare.

Our study has several limitations. First, anesthesia has an uncertain relationship with waking life^54^. Moreover, it remains unclear whether our findings will apply to other non-conscious states, such as sleep and coma^55^. Indeed, our results may not generalize to anesthetics besides propofol. Another limitation is that we did not have enough patients to test for lateralization/dominance. Another limitation is that processes we describe may not be unique to the hippocampus. Indeed, the RNN relies critically on the formatting of its input into binary categories, which must be inherited from other regions. Instead, the goal is to understand how oddball-dependent responses can arise from such input. Furthermore, we cannot definitively conclude that plastic changes in tone identity decoding during the oddball task are a compensatory property of the oddball response, rather than an independent plastic effect such as stimulus-specific adaptation. Finally, our choice to use temporally unpredictable tones may have weakened our sensitivity to tone-related effects as temporally predictable oddballs may maximally drive neural responding^11^.

These results highlight the robust coding of cognitive variables in the hippocampus in the absence of consciousness. Such task-correlated patterns of neural activity are typically thought of as neural correlates of their corresponding cognitive processes, so our observations raise the possibility that those processes may occur in the absence of consciousness. That idea, in turn, suggests that consciousness may have some association with those processes but is not a *sine qua non*. Instead, those processes may occur in a subconscious manner, as for example, we may monitor subconsciously others’ conversations at a cocktail party^56,57^. A good deal of past research has emphasized the central positioning of the hippocampus within the anatomical hierarchy of brain areas; indeed, it may be among the regions most distant from both input and output ends of brain processing^58^. It is therefore generally assumed that the hippocampus will have the most attenuated inputs in the absence of consciousness; our results also invite a reconsideration of that idea.

These results raise the question about what features differentiate conscious from non-conscious processing. Our results, at least, suggest that the key ingredient does not reside in the activity of single units or LFP in the hippocampus. Existing prominent theories consistent with our results include the idea that consciousness involves cross-regional coordination^63–65^, global propagation of local signals^66–68^, or recurrent processing^69–71^. Another possibility is that consciousness is not simply a result of neural activity at a given time but is instead constructed through repeated revisions of current experiences, and that it is this revision process that is impaired^72^.

## METHODS

### Patient recruitment

Experiments were conducted according to protocol guidelines approved by the Institutional Review Board for Baylor College of Medicine and Affiliated Hospitals, Houston TX (H-50885). All recruited patients were diagnosed with drug resistant temporal lobe epilepsy and were scheduled to undergo an anteromesial temporal lobectomy for seizure control. All patients provided written informed consent to participate in the study and were aware that participation was voluntary and would not affect their clinical course. Included patients’ age ranged from 25-54 years old (average 39.6 +/- 11.8), with three females and three male patients. Two resections were on the left side, and three were on the right. In one subject (P4), control recordings were performed in the middle temporal lobe prior to resection to quantify motion. None of the patients reported explicit memory of intraoperative events after the case when discussed in the post-operative care unit or while recovering in the hospital the next day.

Note that we include for comparison purposes a cohort of awake patients listening to podcast stimuli. These patients were recruited from patients undergoing invasive recordings in the epilepsy monitoring unit (EMU) at Baylor St. Luke’s Hospital. Details on methods for this group of patients is found in References 73-77.

### Neuropixels Data Acquisition Setup and Intraoperative Recordings

Neuropixels 1.0-S probes (IMEC) with 384 recording channels (total recording contacts = 960, usable recording contacts = 384) were used for recordings (dimensions: 70μm width, 100μm thickness, 10mm length). The Neuropixels probe, consisting of both the recording shank and the headstage, were individually sterilized with ethylene oxide (Bioseal, CA).^78^ Our intraoperative data acquisition system included a custom-built rig including a PXI chassis affixed with an IMEC/Neuropixels PXIe Acquisition module (PXIe-1071) and National Instruments DAQ (PXI-6133) for acquiring neuronal signals and any other task-relevant analog/digital signals respectively. Our recording rig was certified by the Biomedical Engineering at Baylor St. Luke’s Medical Center, where the intraoperative recording experiments were conducted. A high-performance computer (10-core processor) was used for neural data acquisition using open-source software such as SpikeGLX 3.0 and OpenEphys version 0.6x for data acquisition (AP band (spiking data), band-pass filtered from 0.3k Hz to 10 kHz was acquired at 30 kHz sampling rate; LFP band, band-pass filtered from 0.5 Hz to 500 Hz, was acquired at 2500 Hz sampling rate). We used a “short-map” probe channel configuration for recording, selecting the 384 contacts located along the bottom 1/3 of the recording shank.

Audio was played via a separate computer using pre-generated wav files and captured at 30 kHz or 1,000 kHz on the NIDAQ via a coaxial cable splitter that sent the same signal to speakers adjacent to the patient. MATLAB (MathWorks, Inc.; Natick, MA) in conjunction with a LabJack (LabJack U6; Lakewood, CO) was used to generate a continuous TTL pulse whose width was modulated by the current timestamp and recorded on both the neural and audio datafiles. Online synchronization of the AP and LFP files was performed by the OpenEphys recording software. Offline synchronization of the neural and audio data was performed by calculating a scale and offset factor via a linear regression between the time stamps of the reconstructed TTL pulses and confirmed with visual inspection of the aligned traces.

Acute intraoperative recordings were conducted in brain tissue designated for resection based on purely clinical considerations. The probe was positioned using a ROSA ONE Brain (Zimmer Biomet) robotic arm and lowered into the brain 5-6mm from the ependymal surface using an AlphaOmega. The penetration was monitored via online visualization of the neuronal data and through direct visualization with the operating microscope (Kinevo 900). Reference and ground signals on the Neuropixels probe were acquired separately by connecting to a sterile microneedle placed in the scalp (separate needles inserted at distinct scalp locations for ground and reference respectively).

For all patients (n=6), we conducted neuronal recordings under general anesthesia for at most 30 minutes as per the experimental protocol. All patients were under total intravenous anesthesia (TIVA), with propofol as the main anesthetic per experimental protocol. Inhaled anesthetics were only used for induction and stopped at least an hour prior to recordings. The anesthesiologist titrated the anesthetic drug infusion rates so that the BIS monitor (Medtronic; Minneapolis, MN) value was between 45 and 60 for the duration of the surgical case.^79^ Of note, BIS values range between 0 (completely comatose) and 100 (fully awake), with standard intraoperative values to be between 40 and 60. Specific anesthesia depth was likely flat across the brief time of the experiment. First, recordings took place several hours after anesthesia induction several hours before the end of the procedure, so patients were well into the stable portion of the surgery. Second, the anesthesiologist was maintaining active monitoring and stably controlled anesthesia levels.

For P4, P5, and P6, we recorded neuronal activity during passive auditory stimuli presentation. For P4, a sequence of auditory stimuli (pure tones; f1=1 kHz, f2=3 kHz) were presented with 80-20 probability distribution, with the less frequent auditory stimulus serving as an “auditory oddball stimulus” (n=300 trials). For P5 and P6 we counterbalanced the tones. A sequence of auditory stimuli (pure tones; f1=200Hz, f2=5kHz) were presented with 80-20 probability distribution, while switching the tone frequency designated as the auditory oddball stimulus halfway through (first half, n=150 trials, f2 is auditory oddball; second half, n=150 trials, f1 is auditory oddball). We interleaved a washout period (30 trials) using the same auditory stimuli but presented at 50-50 probability distribution in between the two counterbalanced tasks. The auditory pure tone stimuli were presented for a 100 ms duration, and the intertrial interval for the auditory oddball task was randomly drawn from between 1-3s.

Sound stimuli for the auditory oddball task consisted of high- and low-pitched tones. The low-pitched tone was 100 ms duration and 200 Hz, approximating a square wave. The high-pitched tone was 100 ms duration and 5 kHz frequency, also approximating a square wave. These stimuli were constructed to have distinct perceived pitch and salient onset structure. Stimulus waveforms were matched in amplitude. Sounds were delivered in stereo, using a sound delivery system that was calibrated in the testing suite (B&K type 4939-A-011 calibration mic and NEXUS 4939-A-011 conditioning amplifier). Both speakers had a relatively flat frequency response (+/-5dB) across the utilized frequency range (200-6000Hz) and no high or low frequency roll-off.

For P6, P8, P9, and P11 we also recorded neuronal activity during podcast episodes. P6 listened to three stories, each approximately 7 minutes long, taken from The Moth Radio Hour (https://themoth.org/podcast). The stories were “Wild Women and Dancing Queens”, “My Father’s Hands” and “Juggling and Jesus”. Each episode consists of a single speaker narrating an autobiographical story. P8 listened to “Why We Should NOT Look for Aliens – The Dark Forest”, an educational video created by the Kurzgesagt group (Kurzgesagt GmbH; Munich, Germany) (https://www.youtube.com/watch?v=xAUJYP8tnRE). The selected stories were chosen to be varied, engaging, and linguistically rich.

### Micro CT

Since recordings were only performed in tissue planned for resection, we first removed a small cube of tissue around the probe and then proceeded with the remainder of the resection. The cube specimens were processed following previously described methods.^80^ In brief, resected specimens were fixed in 4% PFA for 16 hours at 4°C. They were then stabilized using a modified Stability buffer (mStability), containing 4% acrylamide (BIO-RAD, cat. No. 1610140), 0.25% w/v VA044 (Wako Chemical, cat. No. 017-19362), 0.05% w/v saponin (MilliporeSigma, cat. No. 84510), and 0.1% sodium azide (MilliporeSigma, cat. No. S2002). Samples were equilibrated in the hydrogel solution for 16 hours at 4°C before undergoing crosslinking at -90 kPa and 37°C for 3 hours. Following crosslinking, excess hydrogel solution was removed, and specimens were washed four times with 1X PBS. Next, samples were immersed in 0.1N iodine and incubated with gentle agitation for 24 hours at room temperature before being embedded in agarose and imaged using a Zeiss Xradia Context micro-CT at 3µm/voxel resolution. The acquired back-projection images were reconstructed using Scout-and-Scan Reconstructor (Carl Zeiss, Ver. 16.8) and converted to NRRD format via Harwell Automated Recon Processor (HARP, Ver. 2.4.1),^81^ an open-source, cross-platform application developed in Python. The 3D volumes were analyzed, and optical sections were captured using 3D Slicer.^82^ All tissue was inspected with a microscope by Dr. Heilbronner and her lab, and no abnormalities were reported.

### Neuronal Data Processing

Because patients were under propofol anesthesia, they did not experience seizures during the surgery, so we did not do any seizure-related data-cleaning. The lack of seizures was confirmed by analysis of the waveform activity by a trained neurologist.

#### Motion-correction

We utilized previously developed and validated motion estimation and interpolation algorithms to correct for the motion artifacts from brain movement.^83^ Motion was estimated via the DREDge software package (Decentralized Registration of Electrophysiology Data software, https://github.com/evarol/DREDge) using either a combination of motion traces obtained using raw LFP and/or AP band data, fine-tuned for individual recordings. Motion-correction was then implemented using interpolation methods (https://github.com/williamunoz/InterpolationAfterDREDge). Both the AP and LFP band data are motion-corrected and utilized for further pre-processing and analysis steps. If the estimated motion led to no improvement in the spike locations then spike sorting proceeded with the motion correction package built into Kilosort 4 without performing interpolation.

#### Unit extraction and classification

Automated spike detection and clustering were performed by Kilosort 2.0 if motion correction was already applied using the DREDge algorithm or KiloSort 4.0^84^ if motion correction was not applied separately. Manually curation of spike clustered was performed using the open-source software Phy.^85^ Unit quality metrics were calculated using SpikeInterface^86^ and were considered single units if they had a d-prime (d’) greater than 1 and fewer than 3% of spikes were violations of a 2ms inter-spike interval refractory period.

#### Local Field Potential data

LFP data was bandpass-filtered between 0.1-20Hz and aligned to task events to extract local ERPs. Gamma band amplitude was calculated in the “high gamma” range, first bandpass filtering it between 70-150Hz and then calculating the absolute value of the Hilbert-transformed complex signal. Given the high correlation between adjacent channels, only 10 channels equally spanning the length of the probe were used to calculate statistics.

### Neuronal Data Analysis

All analyses were performed using custom MATLAB code unless otherwise noted.

#### Motion Analysis

The motion-corrected location estimates were obtained at a 250 Hz sampling frequency using the DREDge algorithm. This signal was downsampled to 10 Hz. The power spectrum of the calculated motion was then estimated using Welch’s overlapped segment averaging estimator for frequencies between 0.1 and 3Hz. The amount of motion was defined as the root mean square error of the location trace of the probes center relative to its average location.

#### Tone Responses

Both single units and multi-units were used for all analyses. A tone responsive neuron was defined as having a statistically significant increase in the average firing rate in the first second after tone onset (shifted by 50ms to account for auditory delay) relative to the preceding 200ms baseline (∝<0.05, Wilcoxon signed-rank test). Visual demonstrations of the peri-stimulus average firing rate were smoothed via a causal Gaussian filter with a standard deviation of 150ms for visualization, however, all statistical analyses were performed on the raw spike count. Response onset latency was computed as the time taken to the peak response. A Mixed Gaussian Model with two components was then fit to the distribution of latencies. Given the trough between the two peaks at 291ms and evidence of average oddball response occurring in the first segment, a window of 0-300ms was used for analysis characterizing tone and oddball selectivity.

#### Neural Tuning

To determine response tuning properties, we modeled trial responses in the peristimulus period using general linear regression models. Neural data in the analysis time window of 0-300ms was used for tuning analyses. Unit response was defined as the average firing rate, LFP power was defined as the root mean square (RMS) value of the bandpass-filtered LFP, and gamma power was defined as the average gamma band amplitude. All vectors were z-scored to allow for comparison of the neural response modulation across units/channels. The independent variables were effects-coded for tone type (frequency 1 vs. frequency 2), trial type (standard vs. oddball), and an interaction term (conjunctive coding). We set the α level at 0.05 to determine if the beta coefficient for the independent variables were statistically significant.

#### Neuronal Population Coding

To determine the information content present in the population, a Support Vector Machine with a linear kernel was trained using 10-fold cross validation for 200 iterations. Accuracy for each iteration was defined as the average accuracy across the 10 folds. Significant coding was defined as the distribution of 200 iterations being statistically different from 0.5 (chance). Algorithm validation was performed by shuffling the dataset and demonstrating that it always performed at chance level. Subsampling was performed to avoid performance bias from an unbalanced dataset (i.e. more standard trials than oddball trials). To investigate the neuronal population response dynamics for tone and oddball encoding as a function of time, we used sets of sequential trials (50 trials) from each of the two counterbalanced blocks (total of 100 trials). For example, the first set was using trials 1:50 and 181:230, whereas the last set was using trials 101:150 and 281:330. Decoding analyses were also run separately for early vs. late trials (first 75 vs. last 75 trials within a 150-trial block) for tone and oddball encoding respectively.

#### Neuronal response learning dynamics

Next, to determine the neural mechanism underlying statistical learning required for oddball detection, we evaluated single-trial response dynamics across the neuronal population. For each trial, we generated a neuronal response population vector. We then computed the Euclidean distance 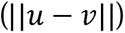 and cosine angle 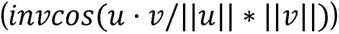 between the mean vector across all standard trials and each individual oddball unit vector, evaluating each as a function of the oddball index.

#### Mixed-effects models

Where applicable, we used mixed effects models to quantify how task conditions affect spike count and other neurophysiological variables while accounting for the hierarchical data structure of multiple subjects, neurons, and channels. For analyses of spike count, we summed spikes over equivalent durations across task conditions and fit a generalized linear mixed effects model (GLME) using a log link function and modeling spike counts as Poisson-distributed. For LFP variables, a linear mixed effects model (LME) was used. All analyses used a random effects structure for neurons or channels nested within-subject.

### Continuous-rate RNN model

We implemented a continuous-rate recurrent neural network (RNN) and trained it to perform an oddball detection task, closely mirroring the one used for the experimental dataset. The network contains 200 recurrently connected units (80% of which are excitatory and 20% of which are inhibitory units). The network is governed by the following equation:

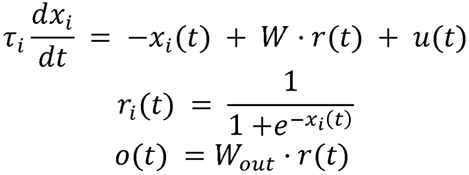

where i represents the synaptic decay time constant for unit i, xi(t) indicates the synaptic current variable for neuron i at time point t, W is the recurrent connectivity matrix (N-by-N; i.e. 200-by-200), and u(t) is the input data given to the network at time point t. u is a 2-by-200 matrix where the first dimension refers to the number of input channels and the second dimension is the total number of time points. A firing rate of a unit was estimated by passing the synaptic current variable (x) through a standard logistic sigmoid function. The output (o) of the network was computed as a linear weighted sum of the entire population of units.

In each trial, the network model receives input data mimicking auditory signals. The input consists of two signal streams, each representing a distinct auditory tone (i.e. tone A vs. tone B; [**Figure 3F,G**]). Only one tone is presented to the network per trial. The model was trained to produce an output signal approaching +1 when Tone A was presented and an output signal approaching -1 when Tone B was presented. To closely replicate the experimental task design, we employed three different sequential contexts during network training. In the first stage, Tone A was presented predominantly (80% of trials), followed by an equal distribution of Tone A and Tone B (50/50) in the second stage. In the third stage, Tone B was predominant (80%).

We optimized the network parameters, including recurrent connectivity, readout weights, and synaptic decay time constants, using gradient descent via backpropagation through time (BPTT). The network was required to achieve over 95% task accuracy in the current context before a new context was introduced. To evaluate the model’s ability to decode both tone identity and oddball context, we performed linear SVM decoding using population activity from the recurrent units. For each decoding analysis, we generated 100 trials for each condition. A linear SVM classifier was trained using 70% of the trials and tested on the remaining 30%, and this procedure was repeated 100 times to estimate decoding accuracy. Separate SVM classifiers were trained for tone identity and oddball context classification.

### Neuronal Data Analysis: Natural Language stimuli

#### Natural language stimuli

All patients were native English speakers. The podcast played during the task was automatically transcribed using Assembly AI (https://www.assemblyai.com/). The transcribed words and corresponding timestamp outputs from Assembly AI were converted to a TextGrid and then loaded into Praat.^87^ The original wav file was also loaded into Praat and the spectrograms and labels and timestamps were manually checked and corrected to ensure the word onset and offset times were accurate. This process was then repeated by a second reviewer to ensure the validity of the time stamps. The TextGrid output of corrected words and timestamps from Praat was converted to a xls and loaded into Matlab and Python for further analysis.

#### Natural Language statistics

Word frequency was defined based on a corpus of movie subtitles spanning a total of 51 million words.^88^ Words that did not elicit a response during the duration of the word were excluded from this analysis. To compare the relative contributions to firing rate, a linear model was trained to estimate the logarithmic firing rate from the logarithmic duration and corpus frequency of each word. Word surprisal values were calculated using the GPT-2 large model^89^ from the Hugging Face Transformers library,^90^ computing the negative log probability of each word conditioned on the preceding context. Specifically, surprisal was defined by the equation: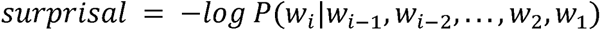 where 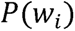 refers to the probability of word *i* given the preceding words.

We utilized the pre-trained fastText Word2Vec model in MATLAB to extract word embeddings for all words in our dataset.^91,92^ This pre-trained model provides 300-dimensional word embedding vectors, trained on 16 billion tokens from Wikipedia, UMBC webbase corpus, and statmt.org, to capture semantic relationships between words. Notably, Word2Vec is a non contextual embedder, so all instances of the same word are represented the same. Some surname words, such as “Harwood” or proper nouns like “Applebee’s” did not have word embeddings and were discarded from the analysis. A simple linear model was trained to predict the firing rate of individual neurons from the semantic matrices using 10-fold cross-validation. Accuracy was defined as the correlation between true and predicted firing rates. Words with 0Hz or above 25Hz were removed from this analysis. To prevent overfitting, Principal Component Analysis (PCA) was used to reduce the dimensionality of the semantic embedding vectors to account for 30% of the variance prior to modeling. This threshold was defined as the minimum of the RMSE of the model that balanced under and overfitting. To predict future or previous words the alignment between words was shifted forwards or backwards, respectively. This relation was then fit with a piecewise exponential decay

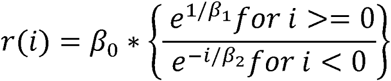

Wherein *β*_0_ is the amplitude of the correlation at 0 lag, and *β*_1_ and *β*_2_ are equivalent to the time constant of the decay for positive and negative lags, respectively.

#### Word embedding, Semantic clustering, and Part of Speech classification

To identify the natural semantic categories present in our word data, all unique words heard by the participants were clustered into groups using a word embedding approach.^93–94^ We used the same 300-dimensional embedding from the prior GLM analysis. To compute and visualize semantic clusters, we first used a t-distributed Stochastic Neighbor Embedding (t-SNE) algorithm on word embedding values to reduce the dimensionality of each unique word based on their cosine distance to all other words, thus reflecting their semantic similarity. Words with similar meanings now have similar 2D coordinates. We then applied the k-means clustering algorithm to these 2D word representations and visualized clustered words on a 2D word map (12 clusters).^95^ We then manually inspected and assigned a distinct label to each semantic cluster and adjusted clusters for accuracy. For example, words bordering the edges of clusters would sometimes get mis-grouped and were manually corrected. The final 12 semantic categories of the words are body parts, places, emotional words, mental words, social words, objects, visual words, numerical words, actions, identity words, function words, and proper nouns. Correction for multiple comparisons was performed using the Benjamini Hochberg approach. A SVM was trained for each semantic category (versus all other categories) using a radial basis function (‘RBF’) kernel. Model training and accuracy metrics were weighted to the relative frequency of each group. We used 10-fold cross validation and 200 iterations.

To extract part-of-speech (POS) for each word in the dataset, we utilized an automated pipeline through Stanford CoreNLP, a natural language processing toolkit.^96^ We initialized a CoreNLPParser with the ‘pos’ tagtype, which specializes in part-of-speech tagging. The transcript was first segmented into sentences based on punctuation. Each sentence was then tokenized and passed through the CoreNLPParser’s tagging function. This process leveraged CoreNLP’s advanced linguistic models to analyze the context and structure of each sentence, assigning appropriate POS tags to individual words. The 15 POS types were: ‘Noun’, ‘Adjective’, ‘Numeral’, ‘Determiner’, ‘Conjunction’, ‘Preposition or Subordinating Conjunction’, ‘Auxiliary’, ‘Possessive Pronoun’, ‘Pronoun’, ‘Adverb’, ‘Particle’, ‘Interjection’, ‘Verb’, ‘Wh-Word’, and ‘Existential’. POS types with fewer than 45 words were removed from analysis. A similar SVM was used for POS.

## STATEMENTS

## Acknowledgements

This project was supported by the Optical Imaging & Vital Microscopy Core at the Baylor College of Medicine and by the McNair Foundation. This project was funded in part by U01 NS121472, the McNair Foundation and the Gordon and Mary Cain Pediatric Neurology Research Foundation.

## Author Contributions

All authors contributed to the design of the work and the acquisition of data KAK, SS, ERC, MCF, JLB, DM, EAM, MM, WM, CWH, ACP, SRH, RK, NR, BYH, and SAS contributed to the analysis and interpretation of data. KAK drafted the manuscript and BYH and SAS substantively revised it. All authors approved the submitted version

## Ethics statement

Experiments were conducted according to protocol guidelines approved by the Institutional Review Board for Baylor College of Medicine and Affiliated Hospitals, Houston TX (H-50885 for the Neuropixel recordings and H-18112 for the EMU recordings).

## Additional Information

Correspondence and requests for materials should be addressed to Sameer A. Sheth. Reprints and permissions information is available at www.nature.com/reprints

## Data availability statement

Core data used for analysis have been uploaded to DABI: https://dabi.loni.usc.edu/dsi/U01NS108923

Code used to implement the computational modeling and analyses in this study is publicly available github: https://github.com/NuttidaLab/rnn_oddball

