## Supplemental Figures for "Plasticity and Language in the Anesthetized Human Hippocampus"

Katlowitz et al., 2025

Supplementary Material

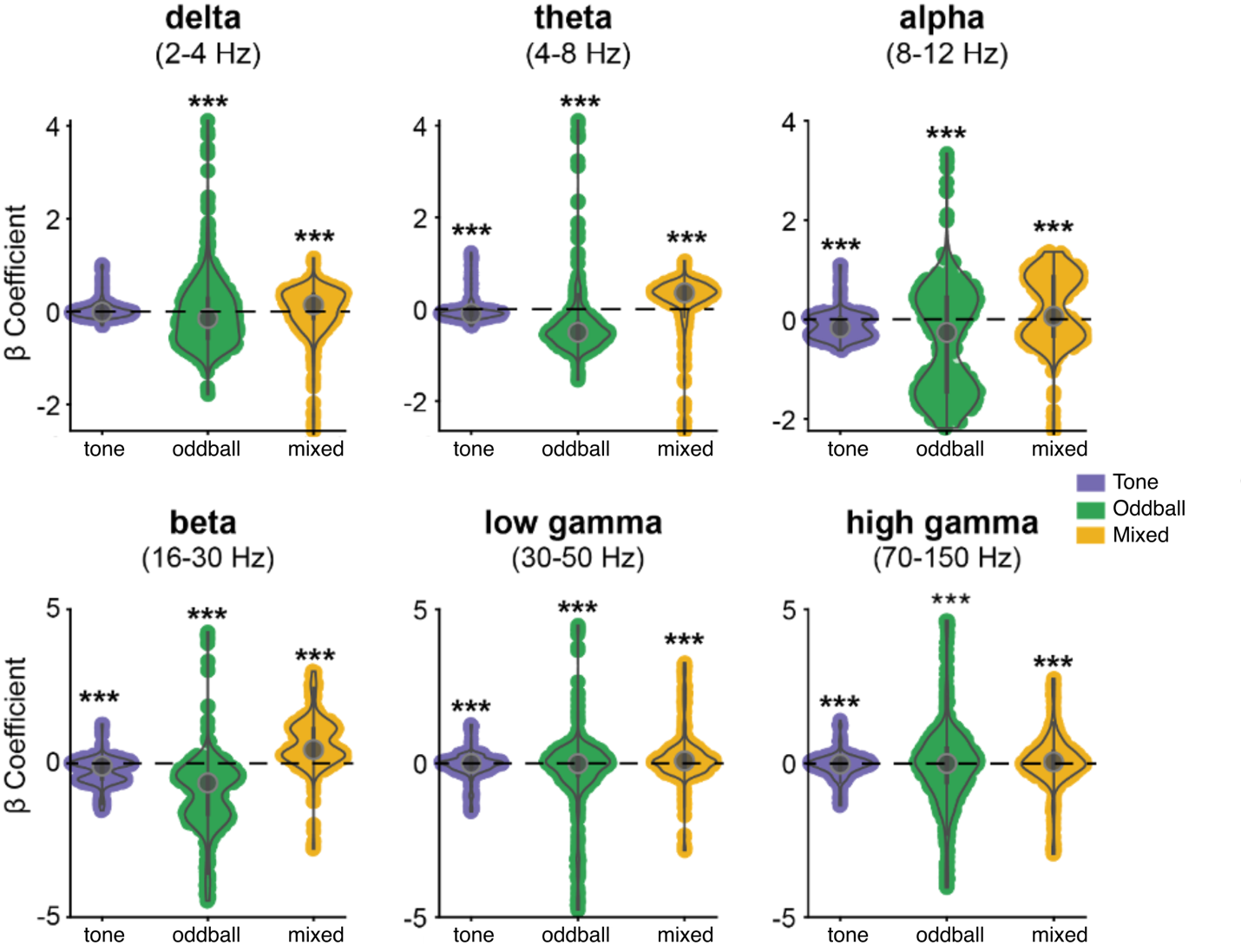

**Figure S1.** Violin plot showing the distribution of β coefficients obtained from a linear regression model run per channel for each frequency band, to determine response modulation as a function of tone identity (tone β, purple, left) oddball identity (oddball β, green, middle), and an interaction/mixed term (mixed β, yellow, right). Asterisks reported statistical significance of the difference between the distribution of individual β coefficients and a distribution with a zero median value (nonparametric Wilcoxon’s sign rank test). *** denotes p-value < 0.0001.

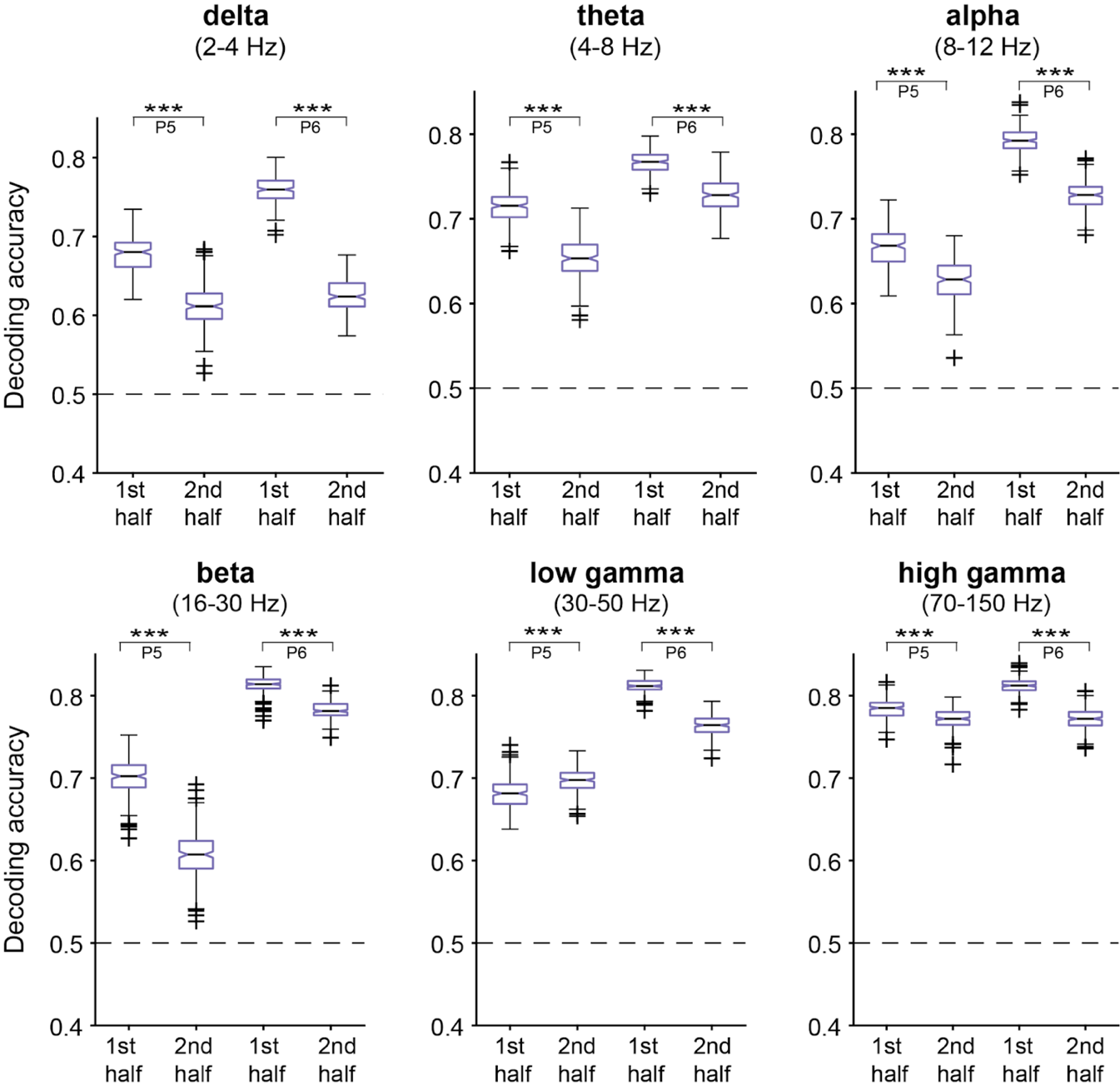
**Figure S2.** Accuracy of tone identity decoding across the population of recorded channels within each pre-defined frequency band for patients P5 and P6, for the first half trials (left) or second half trials (right), combined across both blocks. Statistically significant differences indicated with an asterisk. *** denotes p-value <0.0001

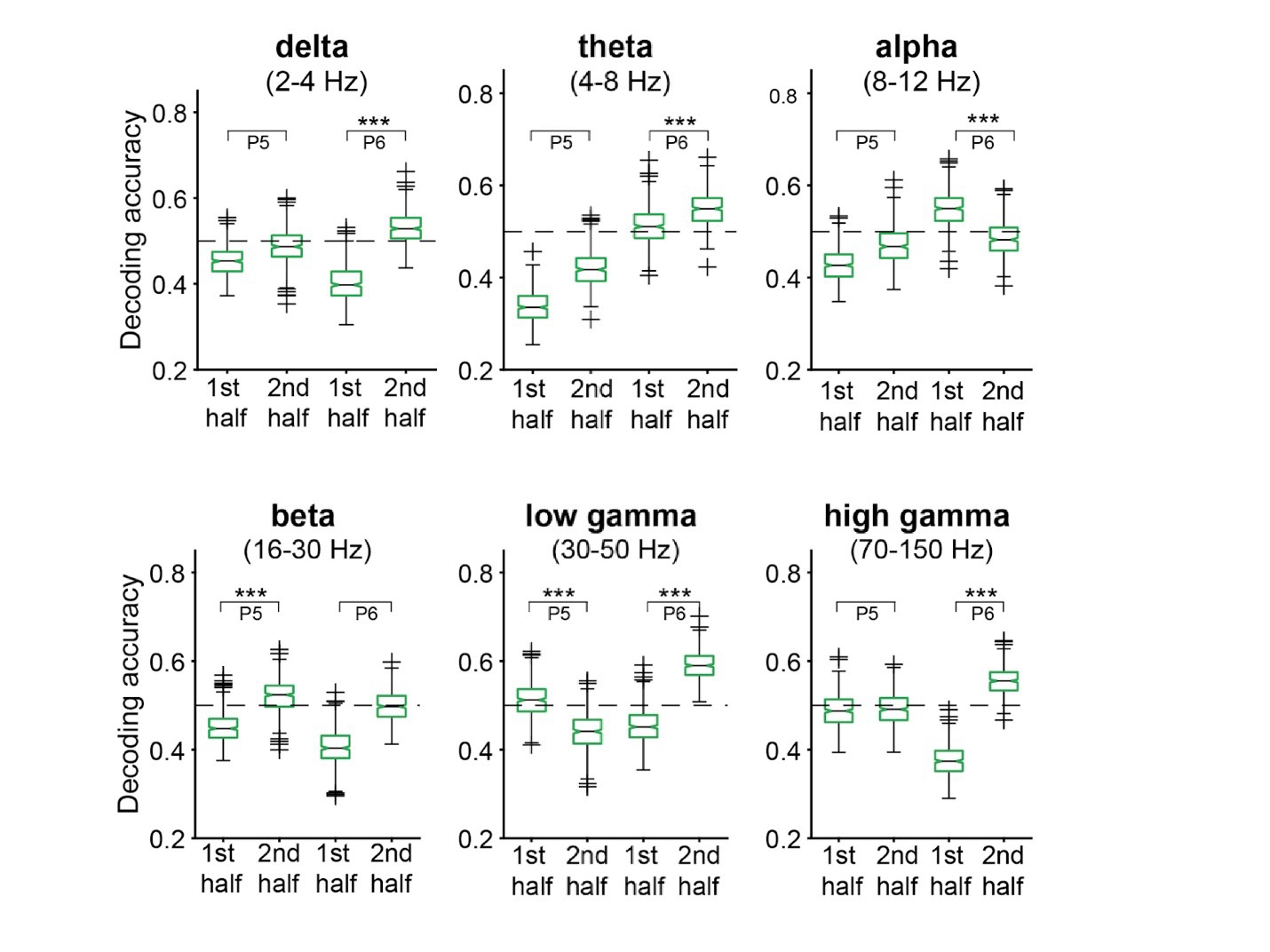
**Figure S3.** Accuracy of oddball identity decoding across the population of recorded channels within each pre-defined frequency band for patients P5 and P6, for the first half trials (left) or second half trials (right), combined across both blocks. Statistically significant differences indicated with an asterisk. *** denotes p-value <0.0001.

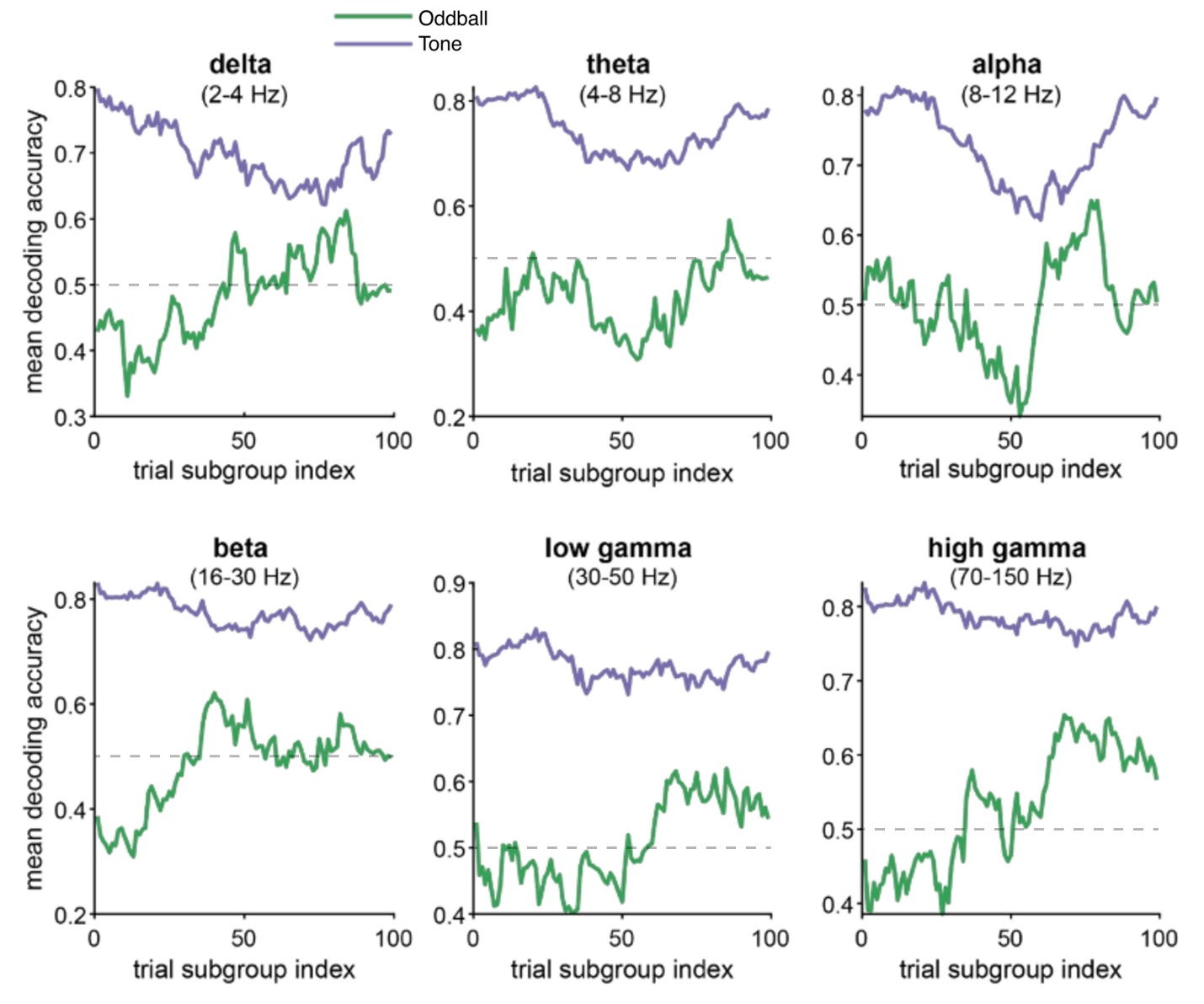
**Figure S4.** Decoding accuracy as a function of trial position for both patients. Each point represents SVM accuracy within a set of 50 trials starting at the index location. Decoding accuracy for tone identity is shown in purple, and for oddball identity in green. Dashed line at 0.5 is chance.

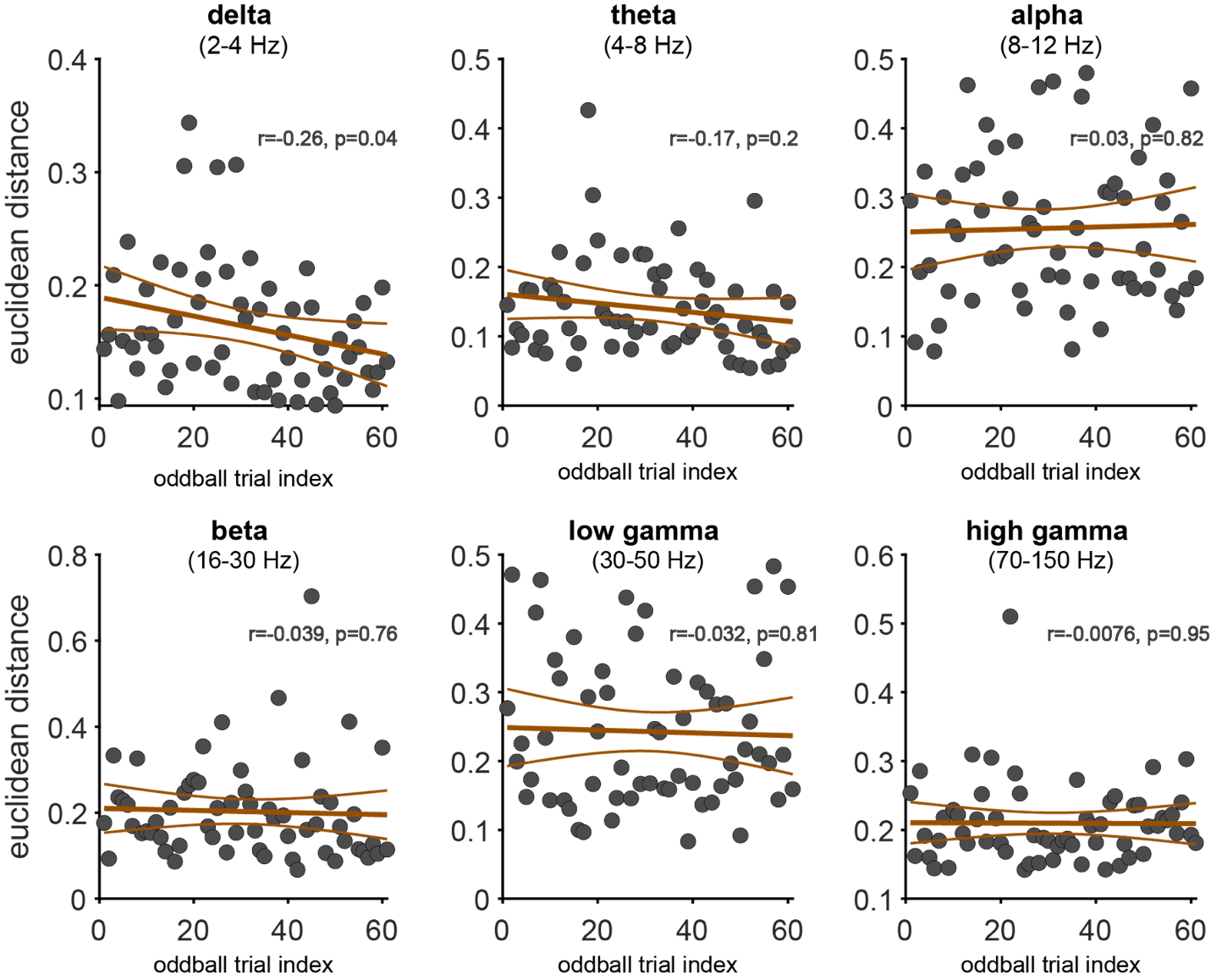
**Figure S5.** Euclidean distance between standard and oddball population response vectors comprising all channels across both patients (n=756 channels), computed for each oddball trial, separately for responses within distinct frequency bands. Each datapoint (in grey) indicates Euclidean distance per trial, and lines show a linear fit with 95% confidence intervals. Analysis of cosine angle (not shown) demonstrated a highly similar trend: for all bands, R was within 0.01 of the equivalent result for Euclidean distance.

**Description of Supplementary Figures 1-5**

Z-scored sensory responses for all channels (computed separately for each frequency band) were modeled as a function of tone identity, context (standard vs. oddball), and their interaction using separate linear regression models. We observed comparable encoding for nearly all terms and for each frequency band (p<0.0001; n=756 channels): i) delta band (2-4 Hz). 2% of channels showed tone encoding; 4% channels showed oddball encoding; 4% channels showed an interaction; ii) theta band (4-8 Hz). 3 % channels showed tone encoding; 3 % channels showed oddball encoding; 3 % channels showed an interaction; iii) alpha band (8-12 Hz). 26% of channels showed tone encoding; 45% showed oddball encoding; 35% showed an interaction; iv) beta band (16-30 Hz). 46% of channels showed tone encoding; 45% showed oddball encoding; 45% showed an interaction; v) low gamma band (30-50 Hz). 12% of channels showed tone encoding; 9% showed oddball encoding; 10% showed an interaction; vi) high gamma band (70-150 Hz). 22% of channels showed tone encoding; 20% showed oddball encoding; 18% showed an interaction. Across the population, the distribution of beta weights was significantly different from a distribution centered at zero median for the following (signrank test on beta values, *** denotes p<0.0001, Figure S1.): i) delta band (2-4 Hz). βtone, p=0.1; βoddball, p <0.0001; βmixed, p<0.0001; ii) theta band (4-8 Hz). βtone, p=0.1; βoddball, p <0.0001; βmixed, p<0.0001; iii) alpha band (8-12 Hz). βtone, p=0.1; βoddball, p <0.0001; βmixed, p<0.0001; iv) beta band (16-30 Hz). βtone, p=0.1; βoddball, p <0.0001; βmixed, p<0.0001; v) low gamma band (30-50 Hz). βtone, p<0.0001; βoddball, p=0.03; βmixed, p<0.0001; vi) high gamma band (70-150 Hz). βtone, p<0.0001; βoddball, p =0.02; βmixed, p=0.002.

Leveraging the power of large-scale recordings, we used a 10-fold cross-validated support vector machine (SVM) to decode stimulus features on a trial-by-trial level across the neuronal population (n=383 channels for p5, n=373 channels for p6). See Figure 2K and the accompanying text for decoding results.

Splitting our task into halves, we found a significant increase in oddball encoding for both patients (paired t-test, p<0.0001 Figure S4). Replicating our findings from the SUA, we also observed a concomitant decrease in tone encoding, again raising the possibility of compensatory mechanisms (p<0.0001, paired t-test for both) (Figure S3).

Using a sliding window of subsets of 50 trials, across most frequency bands, we found a continuous increase in oddball decoding accuracy across the approximately 10-minute duration of the experiment (Figure S4, purple), accompanied by a decrease in tone encoding (p<0.0001, Figure S4, green). Linear fits for evaluating the correlation of the decoding of tone identity and oddball identity as a function of time showed: i) delta band (2-4 Hz). βtone = -0.001, p<0.0001; βoddball=0.00l, p <0.0001; ii) theta band (4-8 Hz). βtone = -6x10^-4, p <0.0001; βoddball=6x10^-4, p <0.0001; iii) alpha band (8-12 Hz). βtone=-5x10^-4, p=0.005; βoddball=4x10^-4, =0.01; iv) beta band (16-30 Hz). βtone=-4x10^-4, p<0.0001; βoddball=0.001, p <0.0001; v) low gamma band (30-50 Hz). βtone= -4x10^-4, p<0.0001; βoddball=0.001, p<0.0001; vi) high gamma band (70-150 Hz). βtone= -3x10^-4, p<0.0001; βoddball=0.002, p <0.0001. These effects are comparable to that observed for single units, further supporting our hypothesis that the neural population was sacrificing its tone responses for the sake of oddball representations over the course of the experiment, suggesting that the hippocampal responses were shifting to represent the salient features of the stimulus.

We created neural vectors of the average standard tone response as well as each individual oddball trial (756-dimensional vectors composed of the mean response of the oddball units). In contrast to our single unit data results, we did not find a statistically significant divergence between standard and oddball vectors over the course of the session (Figure S5).

**Aperiodic slope (oddball)**

Given that multiple LFP bands were sufficient to decode tone and oddball stimuli, we further investigated whether similar results could be obtained using a single aperiodic slope feature. For each trial and channel, we computed the aperiodic slope over the same time interval as other LFP features. We computed the power spectral density between 2 Hz and 40 Hz, performed a log-log transformation, applied a median filter to mitigate the influence of narrow spectral peaks, and computed the linear slope fit (Voytek et al., 2015). Using generalized linear models, aperiodic slope values were modeled as a function of tone identity, oddball status, and their interaction. We found that 12.3% of channels demonstrated significant tone identity encoding, 6.6% of channels demonstrated significant oddball encoding, and 6.1% of channels demonstrated an interaction (p < 0.05 for each channel). Leveraging a 10-fold cross-validated SVM, aperiodic slope reliably predicted tone identity (p5: accuracy = 0.628, p<0.0001; p6: accuracy = 0.677, p<0.0001). Aperiodic slope was insufficient for decoding oddball identity (p>0.05 in both patients).

**Phase-amplitude coupling (oddball)**

After observing successful prediction using both spike and LFP features, we further examined whether stimuli are represented in the temporal relationship between these features in the form of phase-amplitude coupling and spike-frequency coupling. For each trial and channel, we computed the modulation index (Tort et al. 2010) to quantify phase-amplitude coupling of gamma (30-80 Hz) to theta (4-8 Hz) power. Using generalized linear models, modulation index values were modeled as a function of tone identity, oddball status, and their interaction. Phase-amplitude coupling did not distinguish either tone or oddball stimuli: 2.25% of channels demonstrated significant tone encoding, 2.51% of channels demonstrated significant oddball encoding, and 2.78% of channels demonstrated an interaction (p < 0.05). Leveraging SVM decoding, phase-amplitude coupling could weakly predict both tone identity and oddball status in p6 (p6: tone accuracy = 0.505, p < 0.001; oddball accuracy = 0.527, p < 0.005), but not p5 (p > 0.05 for both tone and oddball).

**Spike-frequency coupling (oddball)**

For each cell and trial, we similarly computed the spike-triggered theta power index (STPI) to quantify spike-frequency coupling between each cell’s spike timing and instantaneous theta power. For each trial we computed instantaneous theta power from the LFP channel where spike amplitude for the given cell was measured at highest amplitude, sampled theta power values at the time of each spike and computed the z-score of averaged theta power sampled at spike times relative to the total distribution of theta throughout the trial. Using generalized linear models, STPI values were modeled as a function of tone identity, oddball status, and their interaction. 20% of cells demonstrated significant tone encoding, 14.7% of cells demonstrated significant oddball encoding, and 12.7% of cells demonstrated an interaction (p < 0.05). Using SVM decoding, spike-frequency coupling predicted tone identity in both patients (p5: tone accuracy = 0.589, p < 0.0001; p6: tone accuracy = 0.555, p < 0.0001), but not oddball status (p > 0.05 for both patients).

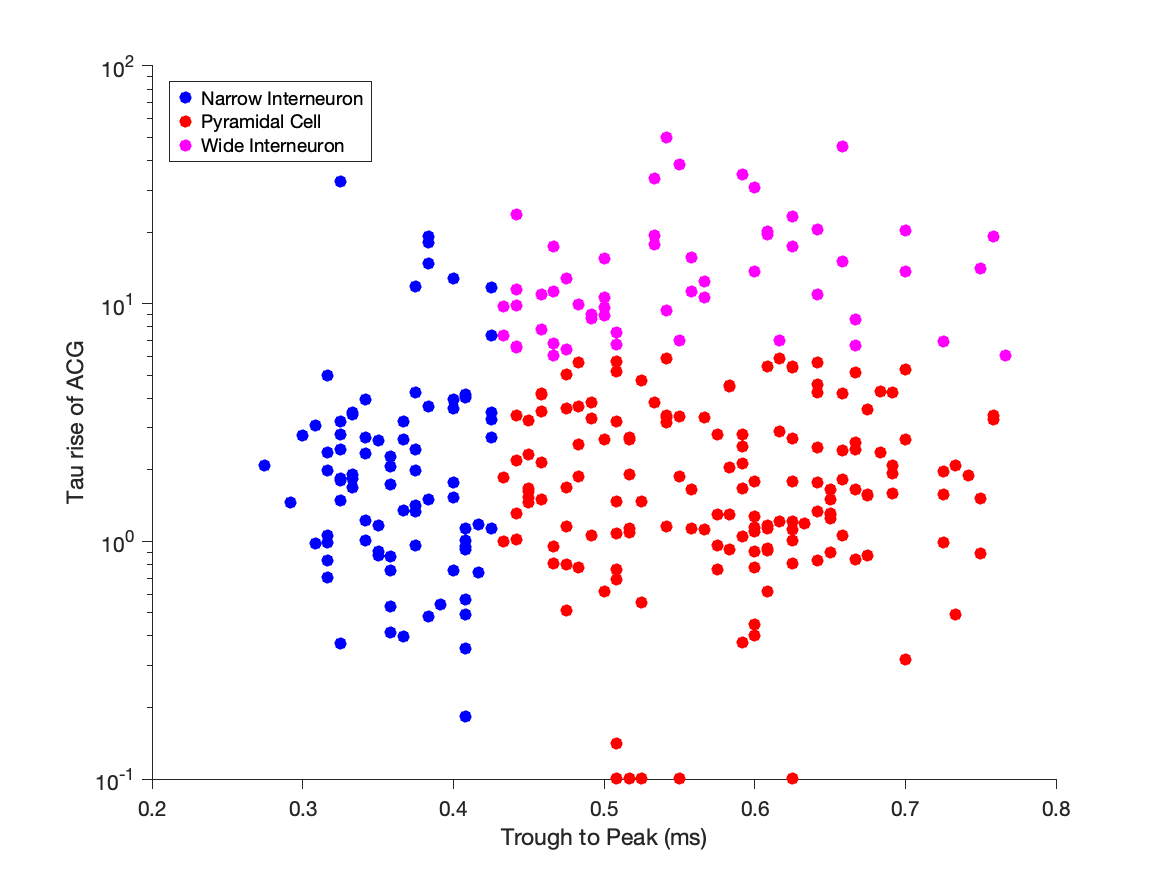

**Figure S6: Categorization of cell types from unit features.** The tone response was significant in a larger proportion of interneurons than pyramidal cells (pyramidal cells: 46.7% responders; interneurons: 73.6% responders; p = 0.0142, chi-squared test). In the tone/oddball experiment (Figs. 2 and 3), we classified 45.3% of cells as pyramidal cells, 51.2% of cells as narrow interneurons, and 3.5% of cells as wide interneurons using conventional analysis tools (CellExplorer) based on their spike waveform and autocorrelogram features. In the language analysis (**Figure 4**), we classified 65.3% of cells as pyramidal cells, 24.6% of cells as narrow interneurons, and 10% of cells as wide interneurons.

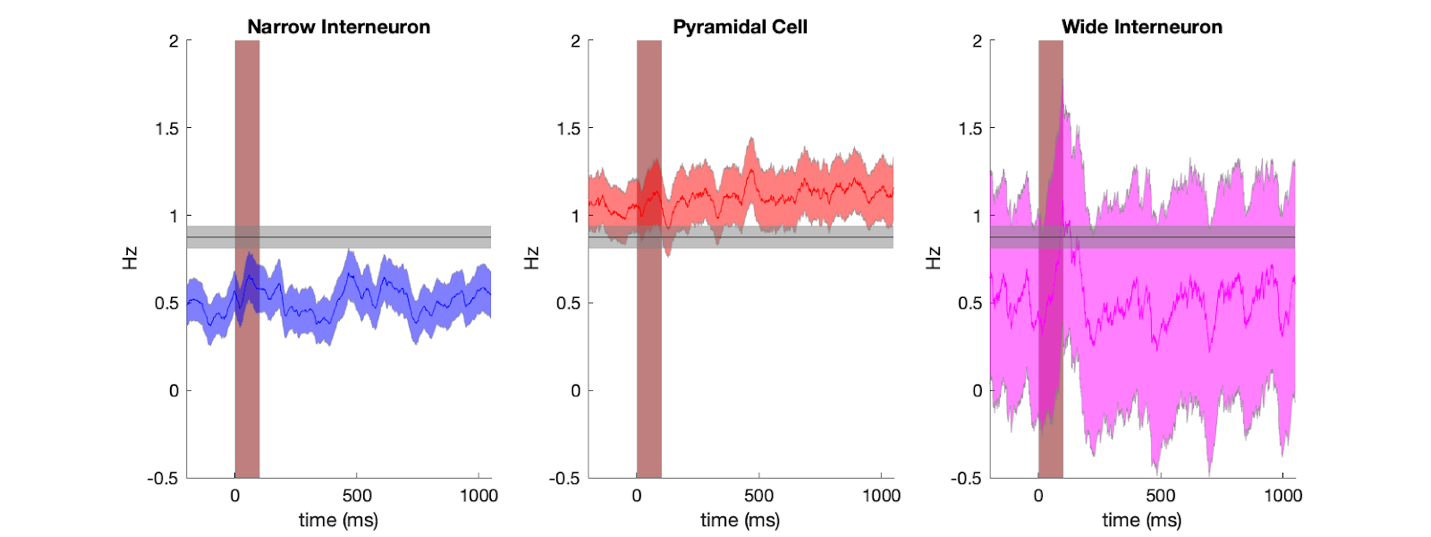

**Figure S7: Tone response by cell type.** Each column plots the firing rate over time (mean ± SEM across cells) in response to tones. Vertical line: duration of the tone response. Horizontal line: baseline firing rate (mean ± SEM across all cells).

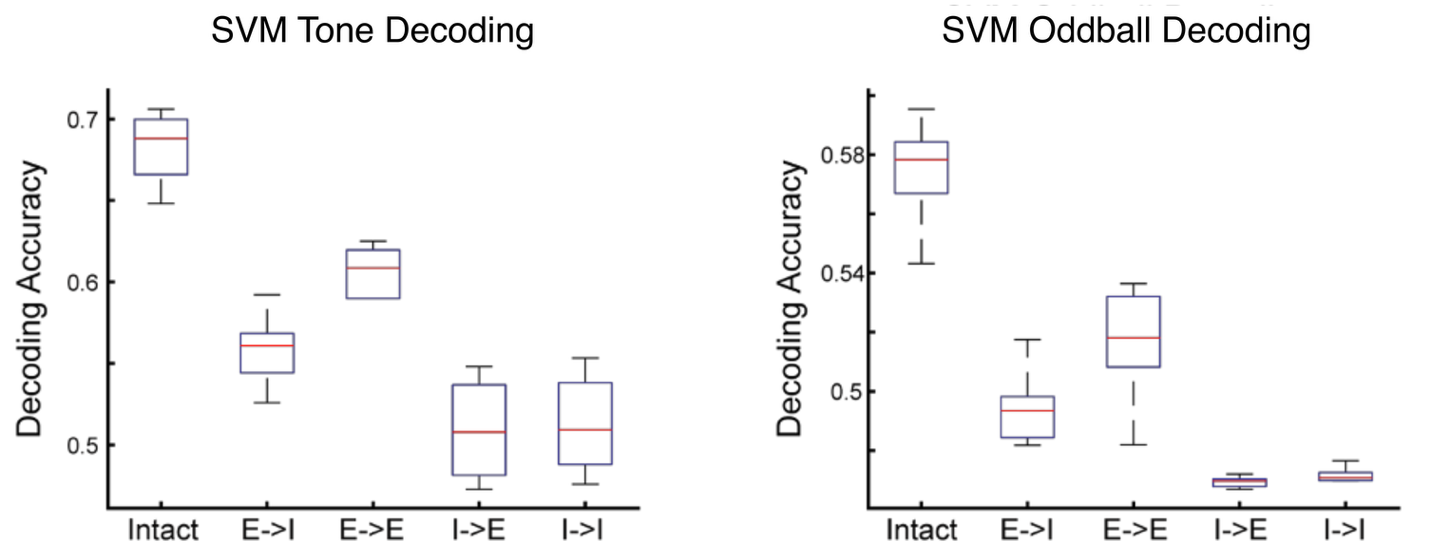

**Figure S8.** Inhibitory connections were important for encoding both tone identity and oddball context. Each subtype of recurrent connection in the trained EI-RNN (E->I, E->E, I->E, and I->I) was lesioned by setting the corresponding weights to zero.

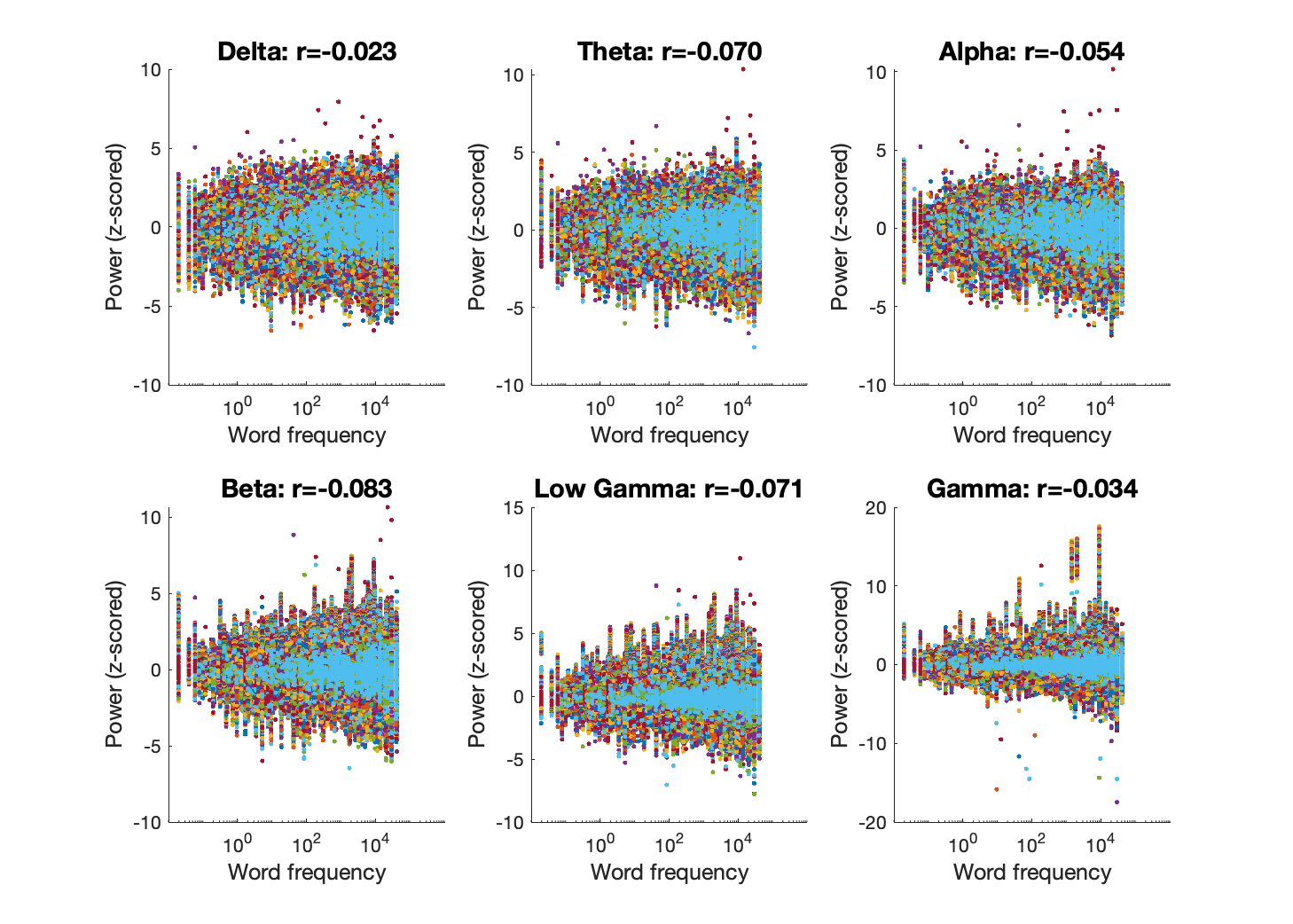

**Figure S9.** Correlation between word frequency and evoked power. Following analysis of the tone experiment (Fig. S1-S6), Z-scored LFP power features were similarly computed for each trial, frequency band, and channel in the 4 patients that participated in the language task. In all 6 bands, power demonstrated a modest but significantly negative correlation with word frequency (p<0.0001; n = 384 channels in each of 4 patients), contrasting the results of single unit analysis.

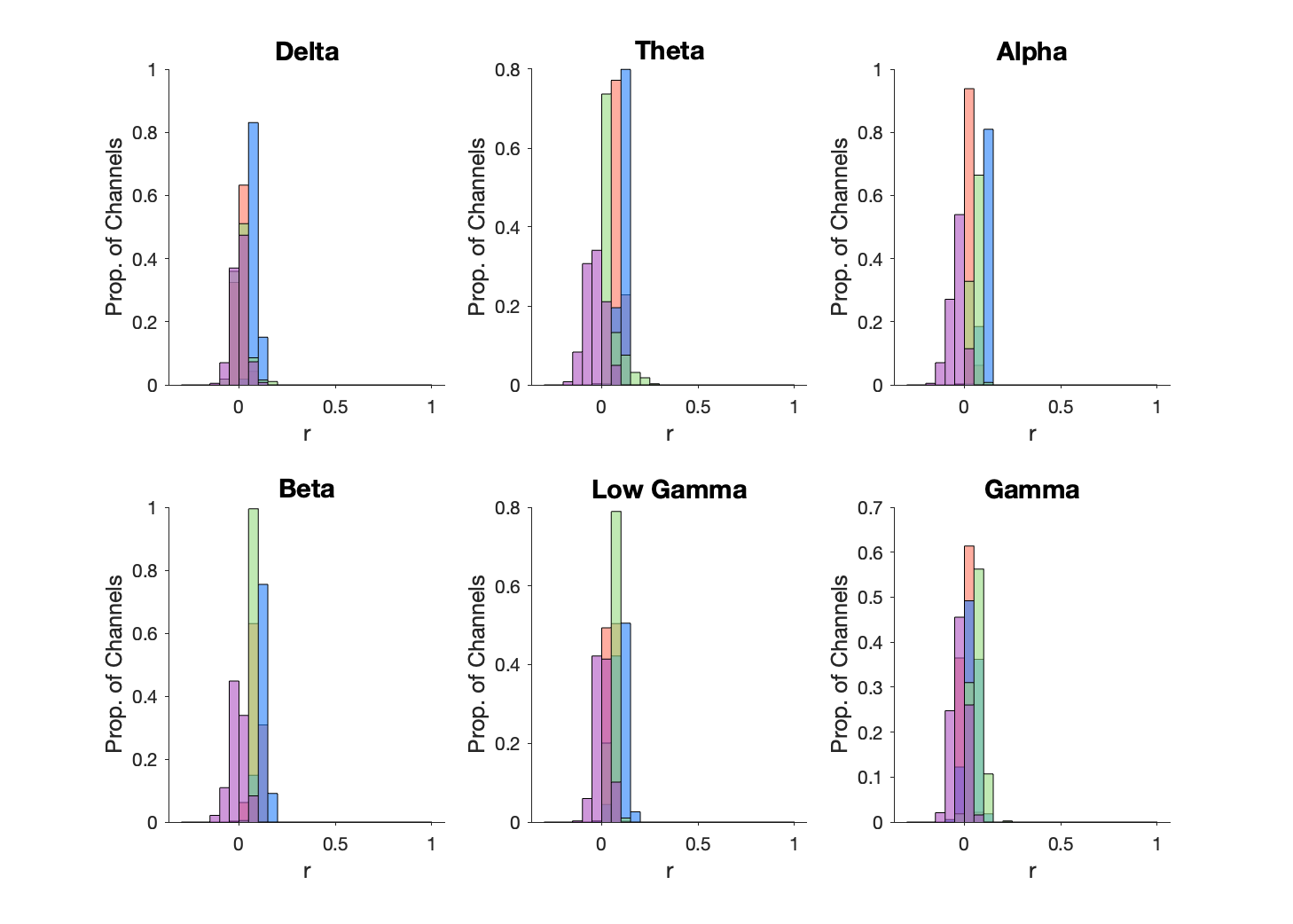

**Figure S10.** Semantic embedding prediction performance by LFP power band. Each panel recreates the analysis of Fig. 4C, showing the average correlation between true power and predicted power of a linear model regressing LFP power in the given band vs. the semantic embedding of each word, computed separately for each channel and grouped by patient. Colors indicate distributions for 4 different patients. For all bands, the RMSE of a linear model significantly outperformed models trained on shuffled data, though with reduced prediction performance relative to single units (p < 0.05; delta: mean R = 0.029, 46% of channels were significant; theta: mean R = 0.054, 75.8% significant channels; alpha: mean R = 0.039, 73.7% significant channels; beta: mean R = 0.067, 76.4% significant channels; low gamma: mean R = 0.052, 76.3% significant channels; gamma: mean R = 0.021, 39.3% significant channels).

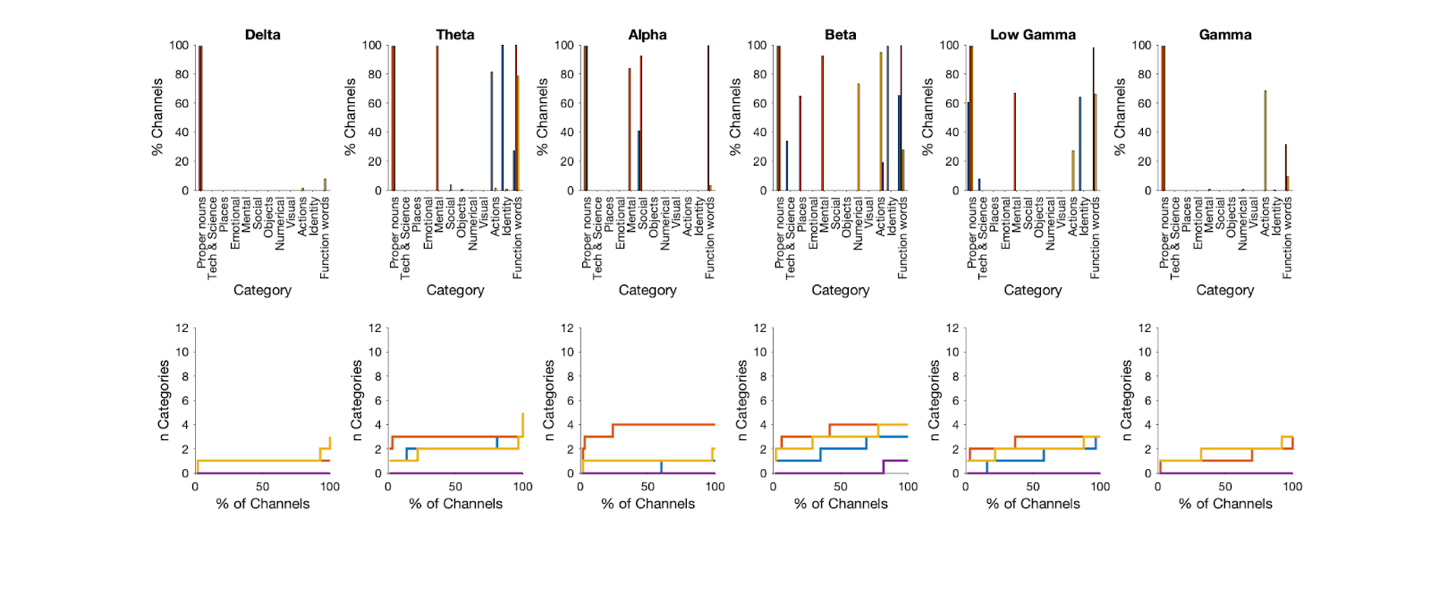

**Figure S11.** Semantic category prediction performance by LFP power band. Each column recreates the analysis of Fig. 4F and Fig. 4G, quantifying the percentage of channels significantly encoding a specific semantic category vs. other categories. Colors indicate distributions for 4 patients. LFP prediction was overall less significant than single units, with higher variability across patients. Beta and low gamma bands were the strongest predictors of semantic category. Overall, semantic category representation was significant but weaker than that of single units. Category discrimination was greatest in the beta band, followed by alpha, theta, and then low gamma. In the delta band, 50% of channels predicted one category and 2% of channels discriminated between 2 categories. In the theta band, 67% discriminated between 2 categories and 31% discriminated between 3 categories. In the alpha band, 60% of channels predicted 1 category, 26% discriminated between 3, and 19% discriminated between 4 categories. In the beta band, 80% of channels predicted 1 category, 50% discriminated between 3 categories, and 21% discriminated between 4. In the low gamma band, 71% of channels predicted 1 category and 20% of channels discriminated between 3 categories. In the gamma band, 50% of channels predicted 1 category and 25% of channels discriminated between 2. For both types of word feature, category discrimination using LFP was more variable across patients and categories than was observed in the single unit analysis. For example, proper nouns were discriminated in 100% of channels in the alpha band for one patient but 0% of channels in another. This observation may reflect the lower-dimensional signal content of LFP recordings, as well as their high signal correlation across densely packed channels of the Neuropixel array.

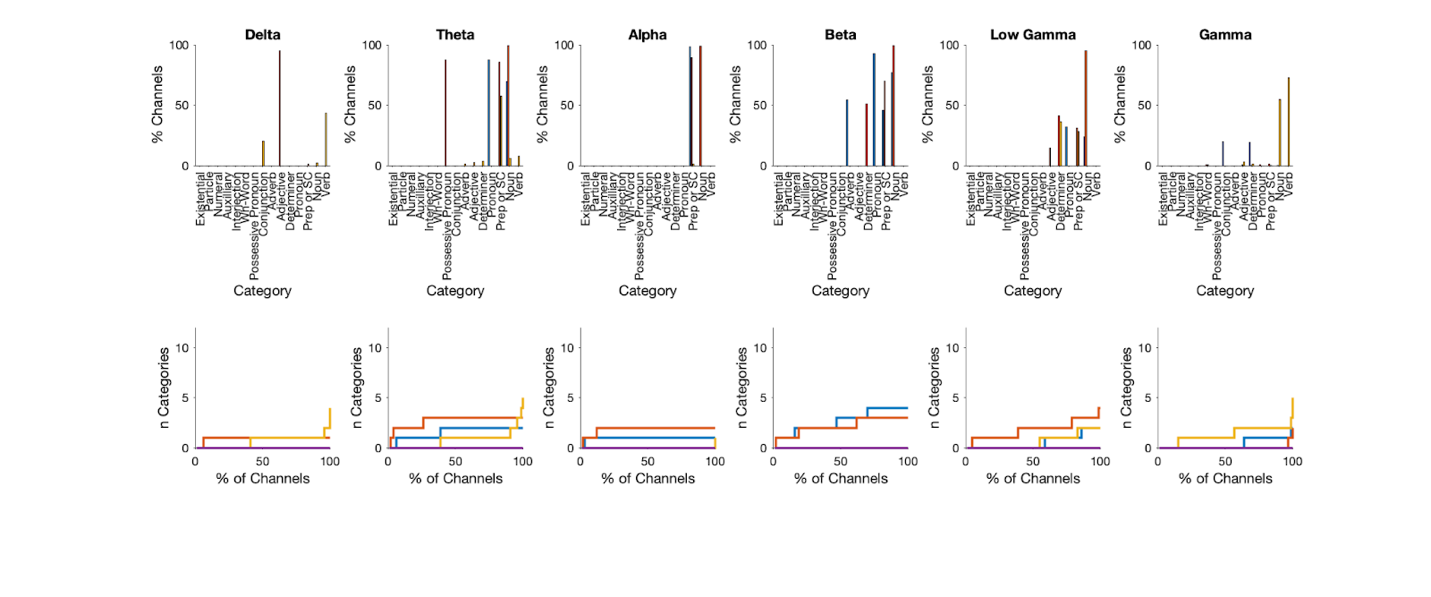

**Figure S12.** Part of speech prediction performance by LFP power band. Each column recreates the analysis of Fig. 4I and Fig. 4J, quantifying the percentage of channels significantly encoding a specific part of speech vs. other categories. Colors indicate distributions for 4 patients. Part of speech discrimination by LFP power was overall less pronounced than semantic category discrimination but demonstrated comparable trends across bands. Part of speech discrimination was greatest in the beta band, followed by theta, then low gamma, and alpha. In the delta band, 39% of channels predicted one category and 1% of channels discriminated between 2 categories. In the theta band, 42% discriminated between 2 categories and 20% discriminated between 3 categories. In the alpha band, 50% of channels predicted 1 category, and 22% discriminated between 2. In the beta band, 42% of channels discriminated between 2 categories, 23% discriminated between 3 categories, and 8% discriminated between 4. In the low gamma band, 46% of cells predicted 1 category, 24% of cells discriminated between 2 categories, and 6% discriminated between 3. In the gamma band, 32% of cells predicted 1 category and 12% of cells discriminated between 2.

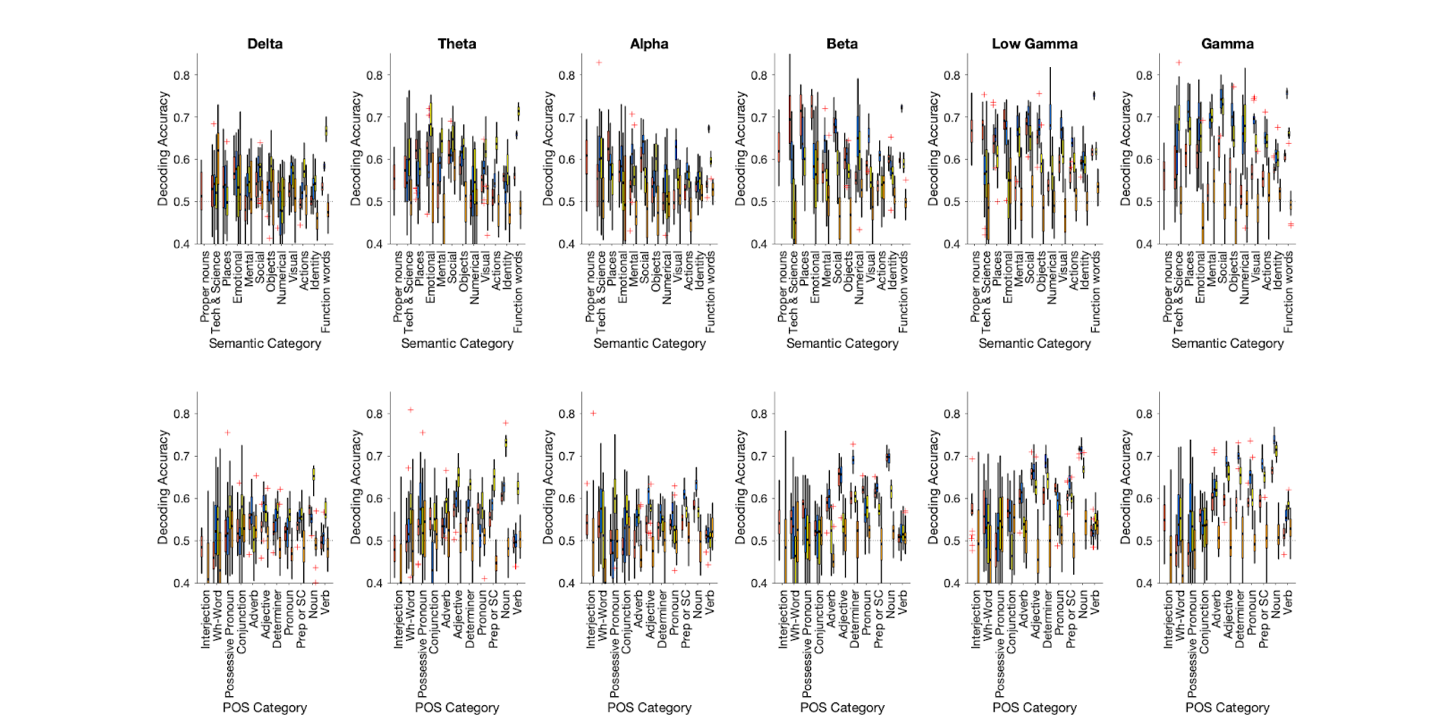

**Figure S13.** SVM classifier prediction performance by LFP power band. Each column recreates the analysis of Fig. 4K (top) and Fig. 4L (bottom), quantifying classifier decoding performance for each word category from bandpower values across channels. Colors indicate distributions for 4 patients. LFP prediction was comparable to that of single units. Frequency bands in the alpha range and above were the strongest predictors for semantic category and part of speech. Decoding accuracy was significantly greater than 0.5 for all bands, but overall lower than that of single units. Semantic category prediction performance was highest for gamma, followed by low gamma and alpha bands (delta: mean accuracy = 0.520, 72.9% significant categories; theta: mean accuracy = 0.536, 81.3% significant categories; alpha: mean accuracy = 0.555, 77.1% significant categories; beta: mean accuracy = 0.517, 85.4% significant categories; low gamma: mean accuracy = 0.564, 85.4% significant categories; gamma, mean accuracy = 0.572, 85.4% significant categories). Part of speech prediction performance was also significant, though modestly lower than that of semantic category prediction, mirroring results of the single unit analysis. Decoding accuracy was highest for alpha power, followed by gamma and low gamma bands (delta: mean accuracy = 0.515, 58.3% significant categories; theta: mean accuracy = 0.523, 60.0% significant categories; alpha: mean accuracy = 0.564, 60.0% significant categories; beta: mean accuracy = 0.473, 68.3% significant categories; low gamma: mean accuracy = 0.535, 70.0% significant categories; gamma, mean accuracy = 0.536, 71.7% significant categories).
